# Cytosolic Ca^2+^ modulates Golgi structure through PKC-mediated GRASP55 phosphorylation

**DOI:** 10.1101/784520

**Authors:** Stephen C. Ireland, Saiprasad Ramnarayanan, Mingzhou Fu, Xiaoyan Zhang, Dabel Emebo, Yanzhuang Wang

## Abstract

It has been well documented that the endoplasmic reticulum (ER) responds to cellular stresses through the unfolded protein response (UPR), but it is unknown how the Golgi responds to similar stresses. In this study, we treated HeLa cells with ER stress inducers, thapsigargin (TG), tunicamycin (Tu) and Dithiothreitol (DTT), and found that only TG treatment caused Golgi fragmentation. TG induced Golgi fragmentation at a low dose and short time when UPR was undetectable, demonstrating that Golgi fragmentation occurs independently of ER stress. Further experiments demonstrated that TG induces Golgi fragmentation through elevated intracellular Ca^2+^ and protein kinase Cα (PKCα) activity, which phosphorylates the Golgi stacking protein GRASP55. Significantly, activation of PKCα with other activating or inflammatory agents, including Phorbol 12-myristate 13-acetate (PMA) and histamine, modulates the Golgi structure in a similar fashion. Hence, our study revealed a novel mechanism through which increased cytosolic Ca^2+^ modulates Golgi structure and function.

## INTRODUCTION

In mammalian cells, the Golgi apparatus is characterized by a multilayer stacked structure of ∼5-7 flattened cisternal membranes, and stacks are often laterally linked to form a ribbon located in the perinuclear region of the cell (*1*). The exact mechanism of Golgi stack formation is not fully understood, but it has been shown that the Golgi re-assembly stacking protein of 55 kDa (GRASP55, also called GORASP2) and its homolog GRASP65 (GORASP1) play essential roles in Golgi stacking (*2, 3*). Both GRASPs are peripheral membrane proteins that share similar domain structures and overlapping functions (*4*). GRASP65 is predominantly concentrated in the *cis* Golgi, whereas GRASP55 is localized on *medial-trans* cisternae. Both GRASPs form *trans*-oligomers through their N-terminal GRASP domains that “glue” adjacent Golgi cisternae together into stacks (*2, 5*) and ribbons (*6–8*). GRASP oligomerization is regulated by phosphorylation; mitotic phosphorylation of GRASP55 and GRASP65 at the C-terminal Serine/ Proline-Rich (SPR) domain inhibits its oligomerization and results in Golgi cisternal unstacking and disassembly (*5, 9*).

The Golgi exhibits different morphology in different cell types and tissues as well as under different conditions. For example, in many secretory cells such as Brunner’s gland of platypus, the Golgi forms large, well-formed stacks (*10*). A plausible hypothesis is that the Golgi adjusts its structure and function in response to different physiological and pathological conditions. For instance, increased neuronal activity causes dispersal of the Golgi at the resolution of light microscopy (*11*), while electron micrographs show reorganization of Golgi membranes in prolactin cells of female rats upon cessation of a sucking stimulus (*12*). In Alzheimer’s disease, the Golgi membranes are dispersed and fragmented in neurons from human brain and mouse models (*13*). Golgi fragmentation is also observed in other neurodegenerative diseases, including Parkinson’s (*14*) and Huntington’s (*15*) diseases and amyotrophic lateral sclerosis (ALS) (*16–18*). In addition, the Golgi has also been shown to be fragmented in lung, prostate and breast cancers (*19–21*). The molecular mechanisms that control Golgi structure and function under disease conditions are so far not well understood.

The Golgi structure can be modulated experimentally such as by molecular manipulations of GRASP55 and GRASP65. Microinjection of antibodies against GRASP55 or GRASP65 into cells inhibits post-mitotic stacking of newly formed Golgi cisternae (*2*). Knockdown (KD, by siRNA) or knockout (KO, by CRISPR/Cas9) of either GRASP reduces the number of cisternae per stack (*22, 23*), whereas simultaneous depletion of both GRASPs causes fragmentation of the entire Golgi stack (*5, 24*). Expression of non-phosphorylatable GRASP65 mutants enhances Golgi stacking in interphase and inhibited Golgi disassembly in mitosis (*23*). Since GRASPs play critical roles in Golgi structure formation, it is reasonable to speculate that physiological and pathological cues may trigger Golgi fragmentation through GRASP55/65 modification, such as phosphorylation (*25*). Using GRASPs as tools to manipulate Golgi stack formation, it has been demonstrated that Golgi cisternal unstacking accelerates protein trafficking, but impairs accurate glycosylation and sorting (*24, 26*). In addition, GRASP depletion also impacts other cellular activities such as cell attachment, migration, growth, and autophagy (*27, 28*).

Protein kinase C (PKC) is a large family of multifunctional serine/threonine kinases that are activated by signals such as increases in the concentration of diacylglycerol (DAG) and/or calcium ions (Ca^2+^) within the cell. In cells, PKCs are mainly cytosolic, but transiently localize to membranes, such as endosomes and Golgi, upon activation (*29, 30*). Membrane association of PKC is via a C1 domain that interacts with DAG present in the membrane. Conventional PKCs (cPKCs) also contain a C2 domain that binds Ca^2+^ ions, which further enhances their membrane association and activity (*31*). Knockdown of atypical PKCs (aPKCs) using siRNA causes a reduction in peripheral ERGIC-53 clusters without affecting the Golgi morphology (*32*). In addition, increased PKC activity has been implicated in cancer (*33, 34*), but the mechanism by which PKC may contribute to invasion, inflammation, tumorigenesis, and metastasis is not fully understood (*35*).

In this study, we performed high-resolution microscopy and biochemistry experiments to determine how the Golgi responds to cellular stresses such as ER stress. While not all ER stress inducers caused Golgi fragmentation, treatment of cells with the Ca^2+^-ATPase inhibitor thapsigargin (TG) resulted in Golgi fragmentation with a low dose and short time in which ER stress was undetectable, indicating that Golgi fragmentation occurs independently of ER stress. Further experiments demonstrated that TG-induced cytosolic Ca^2+^ spikes activate PKC that phosphorylates GRASP55. Interestingly, inflammatory factors such as histamine modulate the Golgi structure through a similar mechanism. Thus, we have uncovered a novel pathway through which cytosolic Ca^2+^ modulates the Golgi structure and function.

## RESULTS

### TG induces Golgi fragmentation and UPR

It has been hypothesized that ER stress and the unfolded protein response (UPR) cause Golgi fragmentation and dysfunction through overloading misfolded proteins into the Golgi (*36*). To test this hypothesis, we performed a time course treatment of HeLa cells with a well-known UPR inducer, TG, which specifically blocks the sarcoendoplasmic reticulum Ca^2+^ transport ATPase (SERCA) (*37*) and causes Ca^2+^ dysregulation (*38*). We assessed the Golgi morphology by co-staining the cells for GM130, a *cis-*Golgi marker, and TGN46, a protein in the *trans*-Golgi network. As shown in Fig. 1A-B, the Golgi became fragmented after TG treatment, and the response was linear over time (Fig. 1A-B; Fig. S1A). Although Golgi fragmentation was more obvious after a longer treatment, it became detectable in shorter treatments such as 10 min. More careful examination of the Golgi morphology by super-resolution fluorescence microscopy demonstrated that the Golgi ribbon was broken down, as the Golgi exhibited as disconnected puncta in the cell. The stacks were also defective, as indicated by the separation of GM130 and TGN46 signals (Fig. 1C-D).

**Figure 1.**
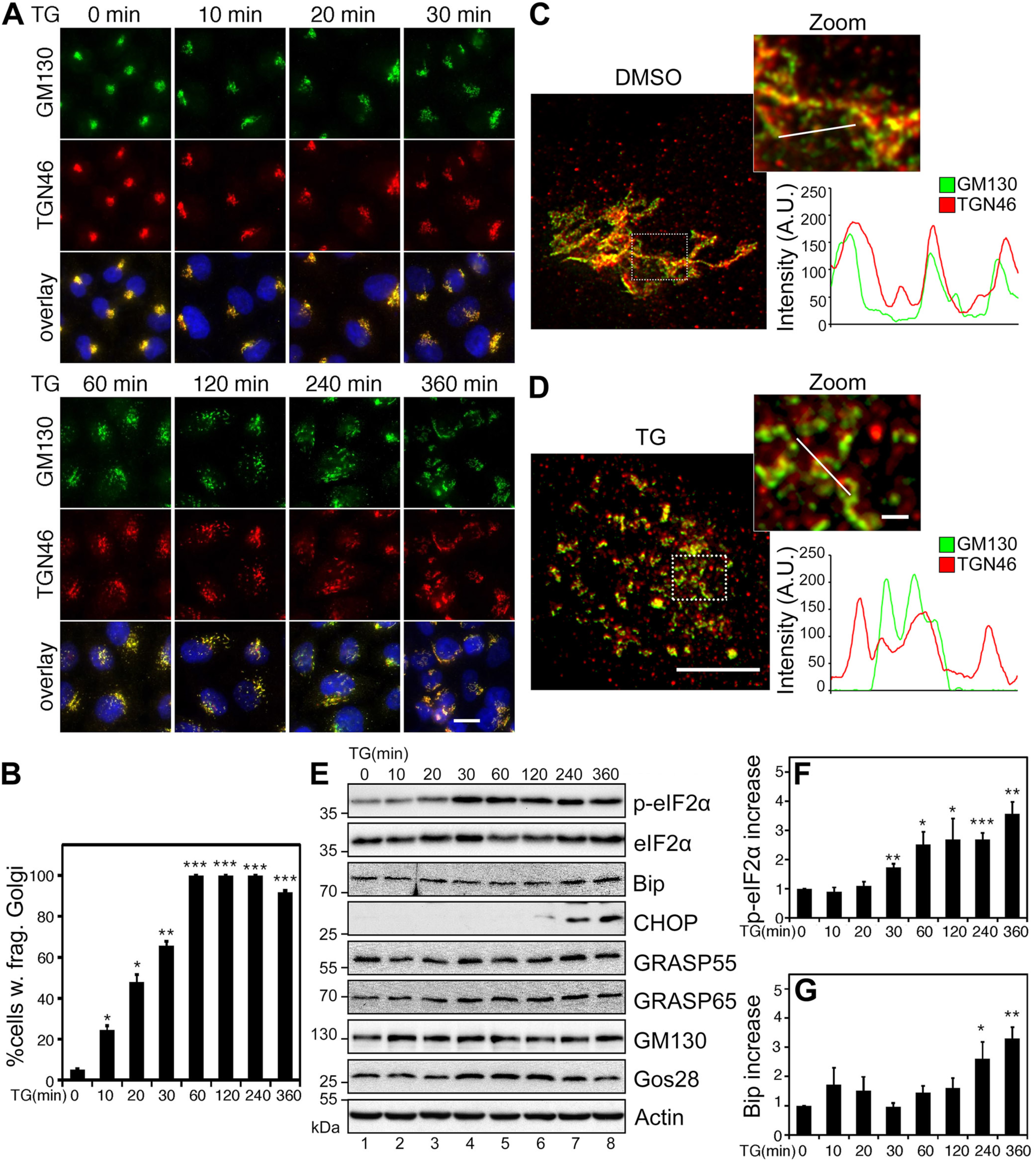
TG induces Golgi fragmentation and UPR. **(A)** Short term TG treatment causes Golgi fragmentation. HeLa cells were treated with 250 nM TG, fixed at the indicated time points, and stained for GM130 (*cis*-Golgi) and TGN46 (*trans*-Golgi). Scale bar, 20 µm. **(B)** Quantitation of (A) for cells with fragmented Golgi using GM130 as the Golgi marker. **(C-D)** Super-resolution images of DMSO (C) and TG-treated (D) HeLa cells. Cells were treated with 2 µM TG for 1 h, stained as in (A) and imaged with a Leica SP8 STED microscope. Indicated areas are enlarged and shown on the right as merged GM130 (green) and TGN46 (red). To quantify Golgi unstacking in these images, relative fluorescence intensity was plotted along a random line through the Golgi region. Note the agreement in peaks in control (C) and relative disagreement in peaks in the TG treated cell (D). Scale bar in main images, 5 µm; in inserts, 1 µm. **(E)** Longer term TG treatment results in ER stress. Cells treated as in (A) were analyzed by Western blot of indicated proteins. Note that TG treatment increases the levels of p-eIF2α, Bip and CHOP. **(F-G)** Quantitation of the ratio of p-eIF2α/eIF2α and the Bip levels from (E), with the no-treatment control normalized to 1. All quantitation results are shown as Mean ± SEM from at least 3 independent experiments; statistical analyses were performed using two-tailed Student’s *t*-tests (*, p ≤ 0.05; **, p ≤ 0.01; ***, p ≤ 0.001).

To correlate Golgi fragmentation with UPR, we performed Western blot of TG-treated cells to assess the levels of several UPR markers, including phosphorylated eukaryotic translation initiation factor 2A eIF2α (p-eIF2α), the ER UPR chaperone binding of immunoglobulin protein (Bip), and the CCAAT-enhancer-binding protein homologous protein (CHOP). As shown in Fig. 1E-G, longer term TG treatment, such as 2 h or longer, caused UPR, as indicated by the increase of all three markers. When the treatment was reduced to 30 min, only the p-eIF2α level increased, while Bip and CHOP did not change. This indicates that the minimal time for UPR to occur is ∼30 min under our experimental conditions. Consistently, no significant increase in the level of any of these UPR markers was detected when the treatment was reduced to below 30 min. Interestingly, the Golgi in a significant proportion of cells was fragmented at this time. Golgi fragmentation was obvious with 10 min TG treatment when UPR was undetectable, and became more prevalent at 30 min treatment (Fig. 1A-B). The fact that Golgi fragmentation occurs earlier than UPR indicates that Golgi fragmentation is unlikely a side effect of ER stress, but rather, occurs independently of UPR.

It is worth mentioning that TG treatment did not affect the level of key Golgi structural proteins, including the Golgi stacking proteins GRASP55 and GRASP65, Golgi tethering proteins GM130 and p115, and the Golgi SNARE Gos28 (Fig. 1E), indicating that TG induces Golgi fragmentation likely through modification rather than degradation of Golgi structural proteins. In addition, TG-induced Golgi fragmentation is reversible; when TG was washed out, the Golgi structure gradually returned to its normal shape (Fig. S1B-D). Consistently, TG-treatment did not induce apoptosis as shown by Annexin V staining. In contrast, staurosporine treatment, which is known to induce apoptosis, increased Annexin V cell surface staining (Fig. S1D-E).

### Tunicamycin (Tu) or Dithiothreitol (DTT) treatment induces UPR but not Golgi fragmentation

To test whether the hypothesis that Golgi fragmentation occurs independently of ER stress applies only to TG treatment or also to other ER stress inducers, we repeated the same set of experiments by treating cells with Tunicamycin (Tu), an antibiotic that induces ER stress by inhibiting N-glycosylation and by the accumulation of misfolded proteins in the ER lumen. 6 h Tu treatment increased the width of the ER cisternae and caused ER fragmentation (Fig. S1F). However, this treatment did not affect the Golgi morphology after 360 min, as indicated by the GM130 and TGN46 signals (Fig. S2A-C). Further analysis of Tu-treated cells by electron microscopy (EM) also did not reveal any significant changes in the Golgi structure (Fig. S2D). The treatment indeed induced UPR, as indicated by the robust increase in the p-eIF2α, Bip and CHOP levels, in particular after 120 min (Fig. S2E-G). Like Tu, Dithiothreitol (DTT) treatment also did not cause Golgi fragmentation, although Bip and CHOP levels increased significantly after 120 min of treatment (Fig. S3A-D). In addition, TG treatment does not seem to affect the organization of the actin and microtubule cytoskeleton (Fig. S3E-F). Taken together, these results indicate that ER stress is unlikely a direct cause of Golgi fragmentation.

### TG induces Golgi fragmentation prior to UPR through elevated cytosolic Ca^2+^

We next sought to decouple the Golgi stress response from UPR after TG treatment. As a complimentary approach to the time-course experiment shown in Fig. 1, we titrated TG (1 - 250 nM) in the treatment. Here we treated cells for 20 min, a time point prior to UPR becoming detectable when cells were treated with 250 nM TG (Fig. 1). The results showed that Golgi fragmentation increased linearly in response to the increasing TG concentration, and importantly, TG at low doses (1 nM to 250 nM) effectively caused Golgi fragmentation (Fig. 2A-B). For comparison, we also assessed UPR in the same cells. As shown in Fig. 2C-E, treatment of cells with up to 250 nM TG for 20 minutes did not cause UPR as indicated by the p-eIF2α and Bip levels.

**Figure 2.**
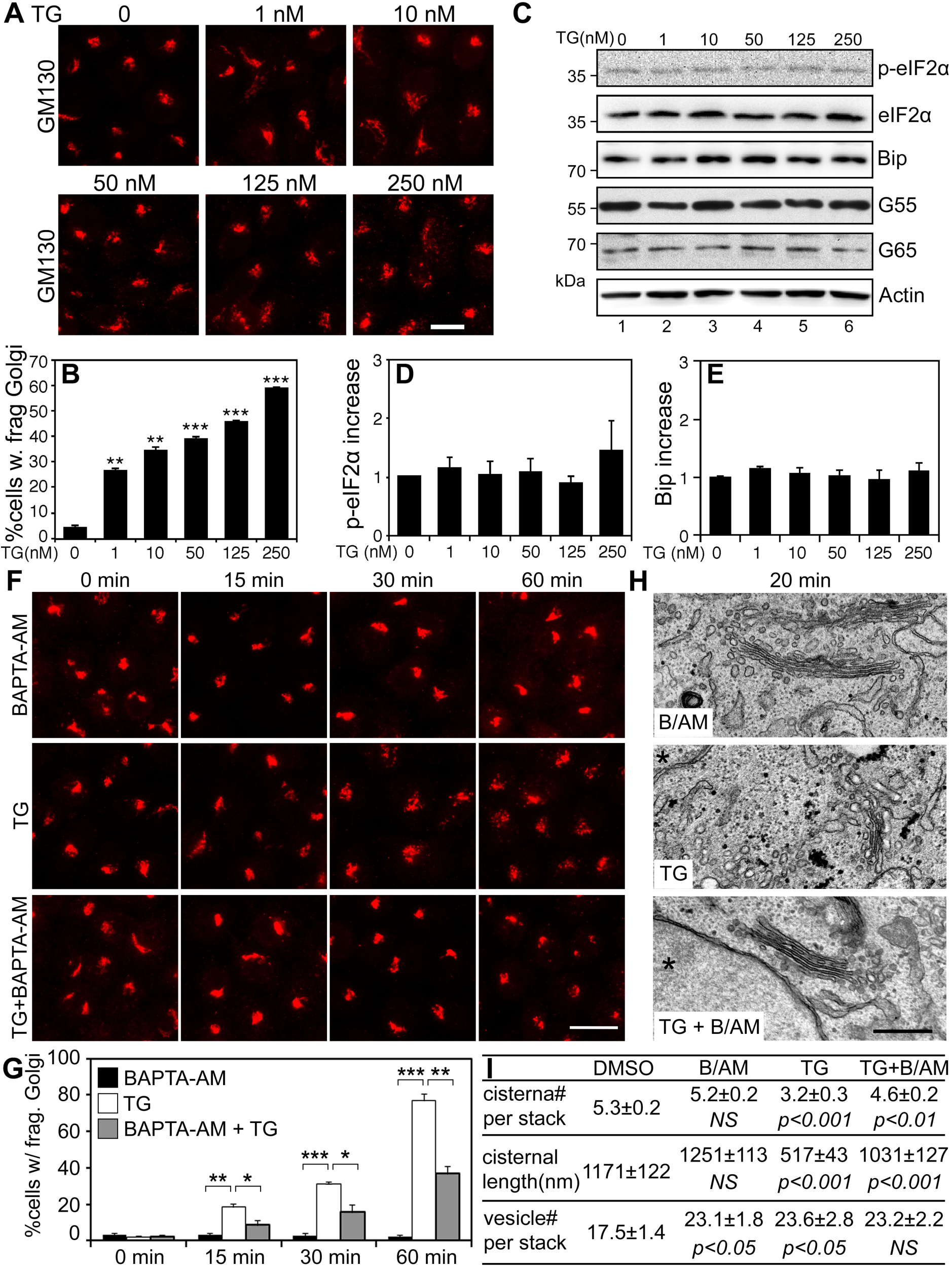
Low concentration of TG induces Golgi fragmentation prior to UPR through elevated cytosolic Ca^2+^. **(A)** HeLa cells were treated with the indicated concentrations of TG for 20 min and stained for GM130. Scale bar, 10 µm. **(B)** Quantitation of cells with fragmented Golgi in (A). **(C)** Cells treated with TG as in (A) were analyzed by Western blots. **(D-E)** Quantitation of p-eIF2α/eIF2α and Bip in (C) from five independent experiments. **(F)** BAPTA-AM inhibits TG-induced Golgi fragmentation. HeLa cells treated with 100 nM TG for indicated times with or without 60 µM BAPTA-AM (B/AM), and stained for GM130. Scale bar, 20 µm. **(G)** Quantitation of (F) from three independent experiments. **(H)** Electron micrographs of Golgi profiles in HeLa cells treated with 250 nM TG and B/AM for 20 min. Note that the Golgi is comprised of bulbous saccules in the TG-treatment, whereas in the B/AM pretreated cells the cisternae appear straight and well-stacked. Asterisks (*) indicate nuclei. Scale bar, 0.5 µm. **(I)** Quantitation of the morphological features of Golgi stacks on the EM images in (H). For statistics, B/AM and TG were compared to DMSO treatment, while TG+B/AM was compared to TG treatment.

We next asked how TG treatment induces Golgi fragmentation. Knowing that TG increases cytosolic Ca^2+^ (*39*), we employed the membrane permeable Ca^2+^ chelater BAPTA-AM to test whether TG induces Golgi fragmentation through cytosolic Ca^2+^. We pre-treated cells with BAPTA-AM alone (60 µM) for 30 min, and then with or without TG (100 nM) for 0, 15, 30, and 60 min (Fig. 2F-G). The result showed that BAPTA-AM significantly prevented TG-induced Golgi fragmentation, while BAPTA-AM alone did not affect the Golgi morphology. Subsequent EM analysis confirmed TG-induced Golgi fragmentation and rescue by BAPTA-AM (Fig. 2H-I). TG treatment reduced the number of cisternae per stack and the length of cisternae, but increased the number of vesicles surrounding each stack. These effects were largely abolished by the addition of BAPTA-AM. These results demonstrated that cytosolic Ca^2+^ is required for TG-induced Golgi fragmentation. Consistent with this notion, treatment of cells with a Ca^2+^ ionophore, ionomycin (Io), also caused Golgi fragmentation (Fig. S4A-B). Therefore, the driving force behind TG-induced Golgi fragmentation is the increased cytosolic Ca^2+^.

### TG-induced Golgi fragmentation increases protein trafficking in the Golgi

As GRASP depletion-mediated Golgi destruction impacts Golgi functions such as protein trafficking (*26, 27*), we examined the effect of TG treatment on the trafficking of the Vesicular Stomatitis Virus Glycoprotein (VSV-G) using the well-established RUSH system (*40*). Cells were transfected with a plasmid that encodes both the invariant chain of the major histocompatibility complex (Ii, an ER protein) fused to core streptavidin and VSV-G fused to streptavidin-binding peptide (SBP). Under growth conditions without biotin, the interaction between streptavidin and SBP retains VSV-G in the ER. Upon the addition of biotin, this interaction is disrupted, resulting in synchronous release of the VSV-G reporter from the ER to the Golgi. Since VSV-G is a glycoprotein, we used endoglycosidase H (EndoH) to distinguish its core (ER and *cis* Golgi) and complex (*trans* Golgi and post-Golgi) glycosylation forms as an indicator of trafficking. As shown in Fig. 3A, TG treatment first slightly decreased VSV-G trafficking at 15 min release, but then increased VSV-G trafficking at 60 and 90 min compared to DMSO control. Our previous studies showed that VSV-G reaches the *cis* Golgi at 15-20 min and *trans* Golgi at ∼90 min (*24, 41*). These results suggest that TG treatment may delay VSV-G release possibly by slowing down its folding; but once it reaches the *cis* Golgi, VSV-G trafficking across the Golgi stack is significantly accelerated. Monensin is known to disrupt the Golgi structure and blocks TGN exit, and thus was used as a control (*42*). As expected, monensin treatment resulted in VSV-G accumulation in the Golgi (Fig. 3A-C).

**Figure 3.**
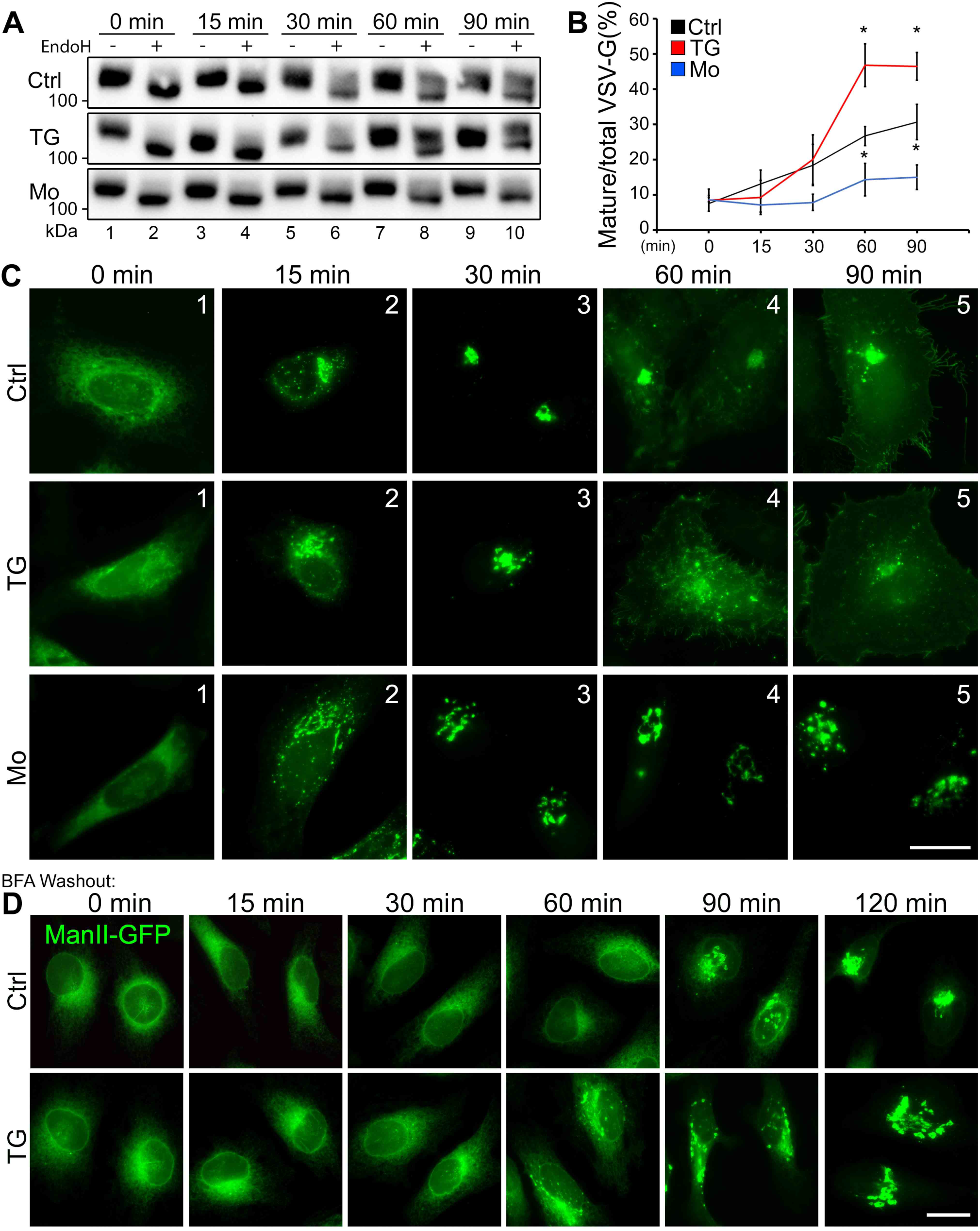
TG-induced Golgi fragmentation has minor effect on protein trafficking. **(A)** Cells were transfected with the Str-li_VSVG wt-SBP-EGFP plasmid for 16 h followed by a 30 min treatment with DMSO, 250 nM TG, or 10 µM monensin (Mo) at 37°C. Cells were then incubated with complete medium containing 40 µM biotin (chase) for the indicated times, lysed and treated with (+) or without (-) EndoH, and analyzed by Western blot for GFP. **(B)** Quantification of (A) for the percentage of EndoH resistant VSV-G from three independent experiments. Quantitation results are shown as Mean ± SEM. Statistical analyses were performed using two-tailed Student’s *t*-tests by comparing to the control (*, p ≤ 0.05). **(C)** Representative images of (A) showing the subcellular localization of VSVG-EGFP at indicated time points after biotin chase. Scale bar, 20 µm. **(D)** Fluorescent images showing the subcellular localization of ManII-GFP in cells treated with DMSO (Ctrl) or 250 nM TG at the indicated time points of BFA washout. ManII-GFP appears in the Golgi area beginning at 60 min, whereas this takes longer in control cells. Scale bar, 20 µm.

To confirm these results using an alternative approach, we treated cells with Brefeldin A (BFA) to accumulate ManII-GFP in the ER. We then washed out BFA and analyzed ManII-GFP in ER-to-Golgi trafficking. The results showed that ManII-GFP started to accumulate in the Golgi at 60 min BFA washout in the presence of TG, while the same observation occurs at 90 min in the control (Fig. 3D).

### TG induces Golgi fragmentation through PKCα activation

Given that phosphorylation of Golgi structural proteins has been shown to cause Golgi fragmentation in physiological conditions such as in mitosis (*2, 5, 23*), as well as in pathological conditions such as in Alzheimer’s disease (*43*), we explored the possibility that phosphorylation of Golgi structural proteins may play a role in TG-induced Golgi fragmentation. We treated cells with staurosporine, a non-selective kinase inhibitor, and a number of specific inhibitors of calcium-related kinases such as protein kinase Cs (PKCs) and Ca^2+^/calmodulin-dependent protein kinases (CAMKs). As shown in Fig. S4C-D, staurosporine significantly reduced Golgi fragmentation in TG-treated cells. In addition, Bisindolylmaleimide I (BIM1), a selective PKC inhibitor, and KN-93, an inhibitor of CAMKII, also partially reduced Golgi fragmentation in TG-treated cells; while the myosin light chain kinase inhibitor ML-7 and the protein kinase A (PKA) inhibitors H-89 and PKI had no such effects (Fig. 4A-B; Fig. S4C-D). These results suggest that either PKC and/or CAMKII is involved in TG-induced Golgi fragmentation. Since both BIM1 and KN-93 inhibitors have pleiotropic effects, we selected two alternative drugs, Gӧ6976 and KN-62, to inhibit PKC and CAMKII, respectively. While Gӧ6976 inhibited TG-induced Golgi fragmentation effectively, KN-62 had no effect (Fig. 4A-B), suggesting a major role of PKC in TG-induced Golgi fragmentation.

**Figure 4.**
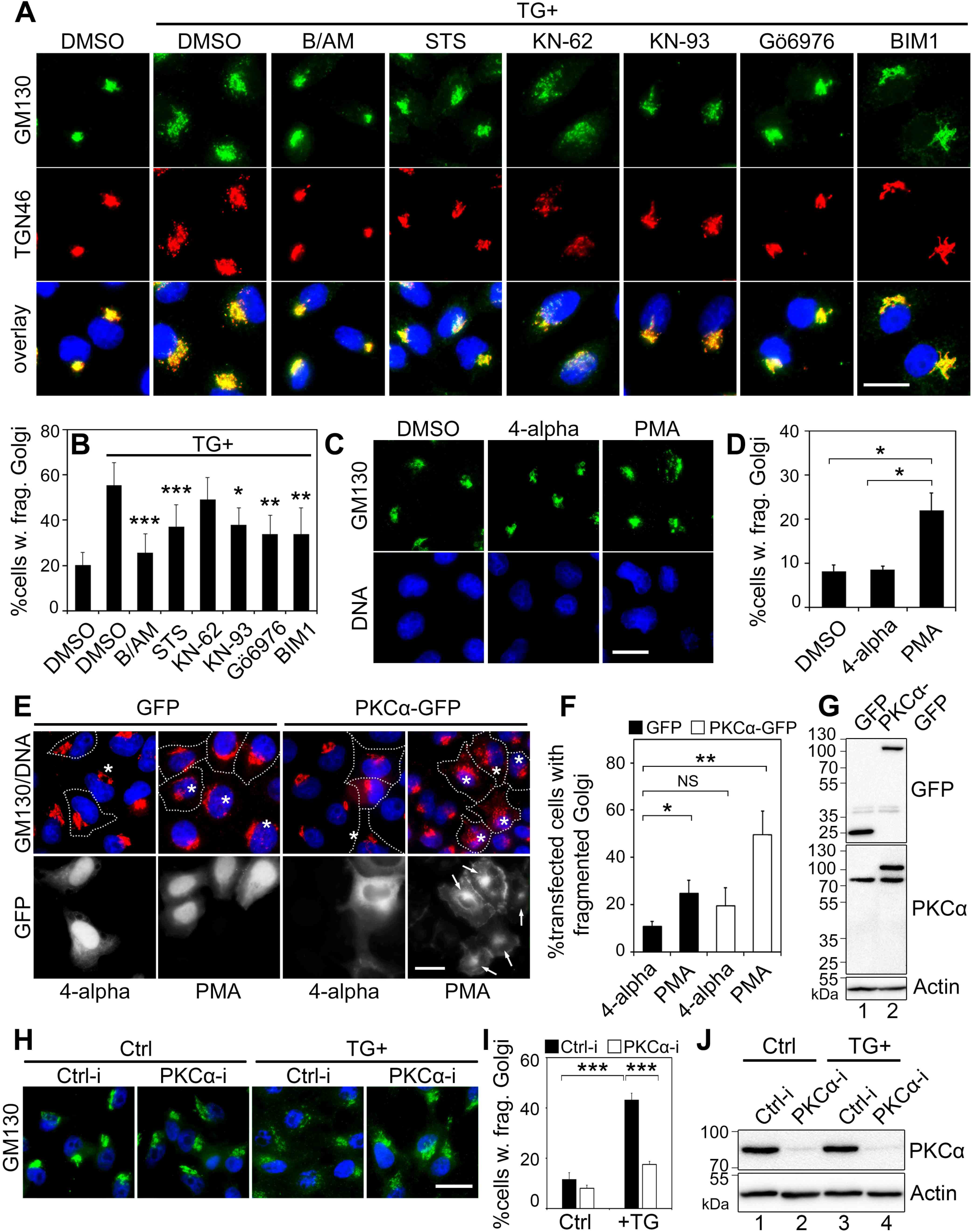
TG induces Golgi fragmentation through PKCα. **(A)** Inhibition of PKC reduces TG-induced Golgi fragmentation. HeLa cells were pre-treated with DMSO, BAPTA-AM (B/AM, 60 µM for 10 min), staurosporine (STS, general kinase inhibitor, 2 µM for 10 min), KN-62 or KN-93 (CAMKII inhibitors, 10 µM and 5 µM, respectively, for 10 min), or BIM1 or Gӧ6976 (PKC inhibitors, 2 µM and 4 µM, respectively, for 10 min), and then with 250 nM TG for 20 min followed by immunostaining of GM130 and TGN46. Scale bar, 20 µm. **(B)** Quantitation of cells in (A) with fragmented Golgi based on the GM130 pattern in the cell. **(C)** HeLa cells were treated with 100 nM PMA for 1 h; 4-alpha-PMA was used as a control. **(D)** Quantitation of cells with fragmented Golgi in (C). **(E)** Activation of ectopically expressed PKCα triggers its Golgi localization (→) and Golgi fragmentation (*). Cells transfected with GFP or PKCα-GFP were treated with 100 nM PMA or 4-alpha for 1 h. Note the fragmented Golgi in these cells upon PMA treatment. Scale bar, 20 µm. **(F)** Quantitation of cells in (E) with fragmented Golgi. **(G)** Western blot of cells from (E) showing that PMA treatment does not affect the PKCα expression level. **(H)** HeLa cells were transfected with control (Ctrl-i) or PKC-specific siRNA for 48 h and then treated with 250 nM TG for 20 min. Cells were fixed and stained for GM130 (green) to show the Golgi structure. Scale bar, 20 µm. **(I)** Quantitation of Golgi fragmentation of cells in (H). All quantitation results are shown as Mean ± SEM from three independent experiments. Statistical analyses were performed using two-tailed Student’s *t*-tests (*, p ≤ 0.05; **, p ≤ 0.01; ***, p ≤ 0.001). **(J)** Cells in (H-I) were blotted for endogenous PKCα to evaluate the siRNA knockdown efficiency.

To further confirm that PKC activation causes Golgi fragmentation, we treated cells with Phorbol 12-myristate 13-acetate (PMA), a widely used PKC activator, and its inactive enantiomer, 4-alpha-phorbol myristate acetate (4-alpha). The results showed that PMA treatment caused Golgi fragmentation, while 4-alpha had no effect (Fig. 4C-D). In addition, expression of CAMKIIβ had no effect on the Golgi morphology (Fig. S4E-F). There have been reports that activation of MAPK/ERK or PKD signaling causes Golgi fragmentation (*44, 45*), we therefore inhibited these two kinases with U0126 or H-89, respectively. Pre-treatment of cells with these kinase inhibitors did not prevent Golgi fragmentation upon the addition of TG (Fig. S4G-H). Taken together, these results indicate that TG induces Golgi fragmentation through PKC activation.

PKC has multiple isoforms including α, βI, βII, γ, δ, ε, η, ζ, and ι (*46*). To identify the PKC isoform responsible for TG-induced Golgi fragmentation, we expressed GFP-tagged PKC isoforms, including all four known classical PKC (cPKC) isozymes (α, βI, βII, γ) that respond to Ca^2+^ stimuli, one from the non-calcium responsive novel PKC (nPKC, δ), and one from the atypical PKC (nPKC, ζ) subfamily (Fig. S5A-B). To enhance the activity of expressed PKC, we also treated cells with PMA, and with 4-alpha as a control. The results showed that expression of PKCα and treatment of cells with PMA increased Golgi fragmentation (Fig. 4E-G; Fig. S5A-B). Interestingly, PKCα-GFP is concentrated on the Golgi upon PMA treatment, as indicated by the colocalization with GM130; while other PKC isoforms did not show the same phenotype (Fig. 4E; Fig. S5B). To further specify that PKCα mediates TG-induced Golgi fragmentation, we knocked down PKCα in cells with siRNA. The results showed that PKCα depletion reduces Golgi fragmentation after TG treatment (Fig. 4H-J). Taken together, these results demonstrate that PKCα activation causes Golgi fragmentation.

### PKCα induces Golgi fragmentation through GRASP55 phosphorylation

Since activated PKCα localizes to the Golgi, we thought it might phosphorylate Golgi structural proteins. To identify potential PKCα targets on the Golgi, we performed gel mobility shift assays on a number of Golgi structural proteins, tethering factors, and SNARE proteins after TG treatment (Fig. S5C). To ensure that the band shift was caused by phosphorylation, we also applied staurosporine (2 µM for 10 min prior to TG treatment) to TG-treated cells to broadly inhibit phosphorylation. Among the proteins tested, GRASP55 and GRASP65 showed a smear above the main bands (Fig. S5C), indicating a partial phosphorylation of the proteins. To increase the resolution of phosphorylated proteins we utilized Phos-tag gels, which showed GRASP55, but not GRASP65, to be significantly shifted up after TG treatment (250 nM, 1 h) (Fig. S5D). TG-induced mobility shift of GRASP55 was not seen upon Tu treatment (Fig. 5A, lanes 2 vs. 3). The shift induced by TG treatment was less dramatic than by nocodazole, which blocks cells from exiting mitosis (Fig. 5A, lanes 3 vs. 4), confirming that TG induced partial phosphorylation of GRASP55. The mobility shift of GRASP55 triggered by TG treatment was abolished by the addition of staurosporine (Fig. 5B, lanes 3 vs. 4; Fig. S5D, lanes 2 vs. 3), validating the mobility shift by phosphorylation. These results demonstrate that TG treatment activates PKCα, which subsequently phosphorylates GRASP55.

**Figure 5.**
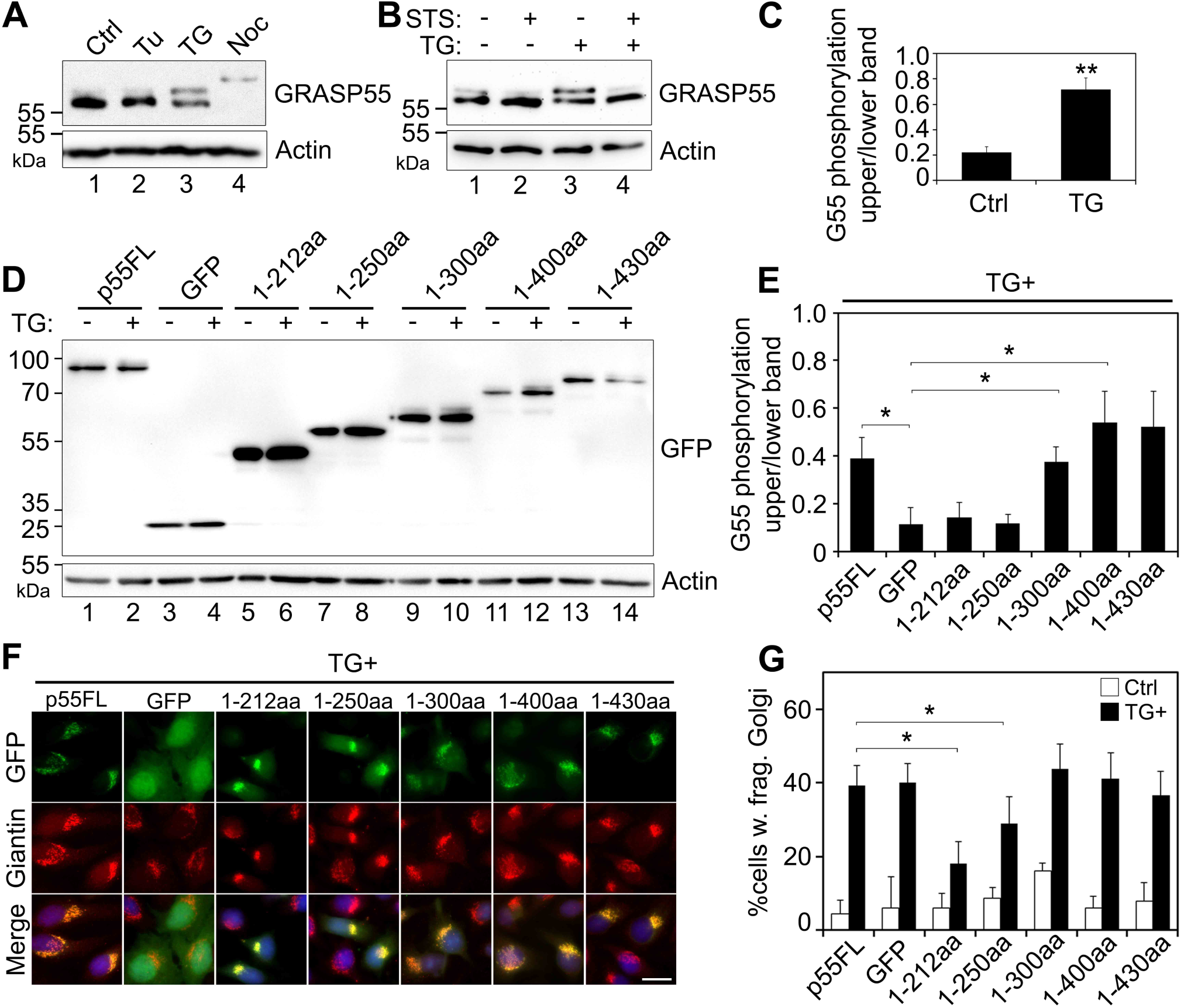
TG induces Golgi fragmentation through GRASP55 phosphorylation. **(A)** GRASP55 is phosphorylated upon TG treatment. HeLa cells treated with Tu, TG or nocodazole (Noc) were analyzed by Western blotting. Note the mobility shift of GRASP55 upon TG treatment compared to control (Ctrl). Nocodazole-arrested mitotic cells were used as a positive control for GRASP55 phosphorylation. **(B)** TG-induced GRASP55 mobility shift is abolished by kinase inhibition. Western blot of cells pre-treated with 2 µM staurosporine (STS) for 10 min and then with 250 nM TG for 1 h. **(C)** Quantitation of GRASP55 phosphorylation in TG treated cells. The intensity of the phosphorylated (upper) band was quantified by densitometry analysis, and plotted relative to the intensities of the full length (lower) band. Shown are the results of relative phosphorylation of GRASP55 from 3 independent experiments. **(D)** Mapping the phosphorylation site on GRASP55 by expressing GRASP55 truncation mutants. Indicated GFP-tagged GRASP55 constructs were expressed in HeLa cells. After TG treatment, GRASP55 was analyzed by mobility shift as in (A). Note the mobility shift in lanes 2, 10, 12, and 14. **(E)** Quantitation of GRASP55 phosphorylation in TG treated cells from (D). **(F)** TG-induced Golgi fragmentation is rescued by the expression of non-phosphorylatable GRASP55 proteins. Cells expressing the indicated GRASP55 constructs were stained for giantin. **(G)** Quantitation of cells in (F) with fragmented Golgi. Results are shown as Mean ± SEM. Statistical analyses were performed using two-tailed Student’s *t*-tests (*, p ≤ 0.05; **, p ≤ 0.01).

GRASP55 contains an N-terminal GRASP domain that forms dimers and oligomers, and a C-terminal serine/proline-rich (SPR) domain with multiple phosphorylation sites (*3, 5*). To map the PKCα phosphorylation site on GRASP55, we expressed GFP-tagged GRASP55 truncation mutants (*47*), treated the cells with TG, and determine their phosphorylation. A visible mobility shift of the GRASP55 variants was observed on the mutants possessing amino acids (aa)251-454, but not the truncated forms shorter than aa250 (Fig. 5D-E). To further determine the functional consequence of GRASP55 phosphorylation, we expressed these constructs and treated cells with TG. The exogenously expressed GRASP55 truncation mutants were targeted to the Golgi, but had no impact on the Golgi structure (Fig. S5E). However, when cells were treated with TG, expression of the N-terminal aa250 or shorter reduced TG-induced Golgi fragmentation; while expression of N-terminal aa300 or longer had no significant effect (Fig. 5F-G). These results demonstrated that phosphorylation of GRASP55 within aa251-300 is important for TG-induced Golgi fragmentation.

### Histamine modulates the Golgi structure through the same pathway as TG treatment

It is known that histamine activates Ca^2+^-dependent PKC isoforms and up-regulates cytokine secretion via the release of calcium from the ER into the cytosol (*48*). It has also been shown that histamine triggers protein secretion and Golgi fragmentation (*49*), but the underlying mechanism has not been revealed. Therefore, we treated HeLa cells with histamine and determined the effect on Golgi morphology. As shown in Fig. 6A-B, histamine treatment induced Golgi fragmentation in a dose and time dependent manner. More than 40% of cells possessed fragmented Golgi after 100 µM histamine treatment for 1 h, a concentration and time often used in previous studies (*50, 51*). Subsequent EM analysis confirmed that histamine treatment induced alterations in the Golgi structure, including fewer cisternae per stack, shorter cisternae, and an increased number of Golgi-associated vesicles (Fig. 6C-D; Fig. S6A).

**Figure 6.**
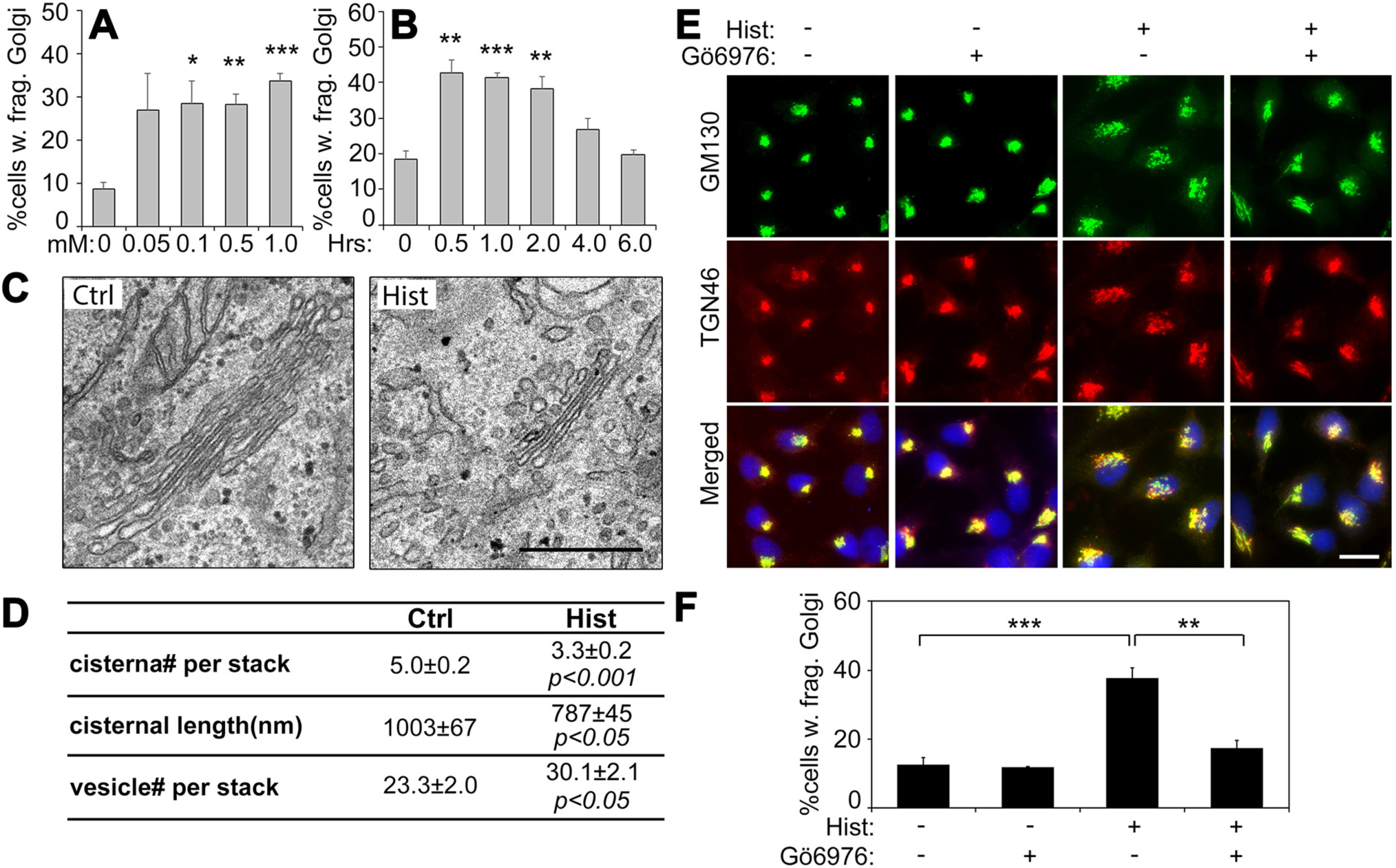
Histamine induces Golgi fragmentation through PKC. **(A)** HeLa cells were treated with indicated concentrations of histamine for 1 h, stained for GM130, and quantified for the percentage of cells with fragmented Golgi. Shown are the quantitation results. **(B)** HeLa cells were treated with 100 µM histamine for the indicated times and analyzed as in (A). Note that histamine induced Golgi fragmentation within 2 h. **(C)** Electron micrographs of Golgi profiles in HeLa cells treated with DMSO control (Ctrl, left panel) or 100 µM histamine for 1 h (right panel). Note the reduced size of the Golgi in histamine-treated cells. Scale bar, 0.5 µm. **(D)** Quantitation of the morphological features of the Golgi stacks in (C). For statistics, histamine treated cells were compared to control cells. **(E)** Cells were pre-treated with (+) or without (-) the PKC inhibitor Gӧ6976 at 37°C for 10 minutes followed by the addition of 100 µM histamine for 60 min. Scale bar, 20 µm. **(F)** Quantitation of cells with fragmented Golgi in (E). Results are shown as Mean ± SEM. Statistical analyses were performed using two-tailed Student’s *t*-tests (*, p ≤ 0.05; **, p ≤ 0.01; ***, p ≤ 0.001).

It has been previously shown that histamine activates Gβγ, which causes TGN fragmentation (*49*). Since HeLa cells do not express Gβγ, and histamine treatment triggers fragmentation of the entire Golgi stack (Fig. 6C-D), the Golgi fragmentation observed in our study may occur through a different mechanism. Indeed, like TG, histamine-induced Golgi fragmentation also depended on PKC, as the addition of the PKC inhibitor Gӧ6976 reduced histamine-induced Golgi fragmentation (Fig. 6E-F). To investigate the functional consequence of histamine treatment on Golgi function, VSV-G trafficking experiments were performed as above. As shown in Fig. S6B-C, histamine treatments slightly decreased VSV-G trafficking within 45 min release, but then began to increase the trafficking speed in the Golgi at later time points. These results demonstrate that histamine treatment impacts Golgi structure and function through a similar mechanism as TG.

## DISCUSSION

In this study, by comparing Golgi fragmentation with ER stress in response to TG, Tu and DTT treatments, we uncovered a novel signaling pathway through which increased cytosolic Ca^2+^ triggers Golgi fragmentation through PKC activation and GRASP55 phosphorylation. Significantly, we also demonstrated that histamine modulates the Golgi structure and function via a similar mechanism, which opens a new window through which we can understand the effect of histamine on cell physiology.

One possible model of Golgi stress is that the expanding capacity of the ER during cellular stress leads to the failure of the Golgi as it is over-burdened with misfolded or improperly folded proteins, affecting its functions like glycosylation (*36*). However, our results do not support this hypothesis for two reasons: First, although all three ER stress inducers, TG, Tu and DTT, all induce ER stress, only TG treatment causes Golgi fragmentation. Second, TG induces Golgi fragmentation at a low dose and time when UPR is undetectable. These results demonstrate that Golgi fragmentation occurs independent of ER stress; instead, the Golgi may possess its own mechanism to sense and respond to stress. Furthermore, our study revealed a novel mechanism that coordinates Golgi structure and perhaps function: TG treatment increases cytoplasmic Ca^2+^, which activates PKC and subsequent phosphorylation of GRASP55, impairing its function in Golgi structure formation. GRASP55 therefore provides the conceptual link between an extracellular cue and Golgi morphological change during stress.

GRASP55 is comprised of an N-terminal GRASP domain (aa1-212) that forms dimers and oligomers and functions as a membrane tether to maintain an intact Golgi structure, and an SPR domain (aa212-454) that undergoes post-translational modifications and functions as the regulatory domain of the protein (*3, 5*). Originally GRASP55 was found to be phosphorylated by ERK2 at T225 and T222 (*44*). Subsequently, additional sites, such as S245 and T249, were identified to be phosphorylated in mitosis, which is required for mitotic Golgi disassembly (*5*). Recently, GRASP55 was discovered to be de-O-GlcNAcylated upon energy deprivation and regulates autophagosome maturation (*47*). These results indicate that GRASP55 is an excellent candidate to function as both a sensor and effector of cellular stresses. Thus, GRASP55 is likely a master regulator of Golgi structure formation, function, and stress responses.

Using GRASP55 truncation mutants we mapped the sites to the aa251-300 region in PKC-mediated phosphorylation. Expression of truncation mutants of GRASP55 that lack this region significantly reduced TG-induced Golgi fragmentation. Previously, it has been shown by mass spectrometry that GRASP55 is phosphorylated on S441 after TG treatment, but the kinase mediating this phosphorylation is unknown (*52*). Although our results are consistent with this previous study, the exact phosphorylation site(s) need further investigation.

Histamine is a neuroendocrine hormone involved in the regulation of stomach acid secretion, brain function and immune response; many of these functions involve secretion (*50, 51, 53*). The role of histamine in immune response is often through the activation of the downstream PKCα. For example, histamine enhances the secretion of granulocyte-macrophage colony stimulation factor (GM-CSF) and nerve growth factor (NGF) in different cell types, both through a PKCα-dependent mechanism (*54*). Interestingly, histamine promotes HeLa cell proliferation and growth, and has been shown to be elevated in cancers where Golgi is fragmented and secretion is enhanced. In our experiments, histamine induced a clear Golgi fragmentation phenotype, confirming a link between histamine and Golgi fragmentation. Additionally, expression of PKCα, but not other PKC isoforms, along with a stimulation with PMA, exhibited an additive Golgi fragmentation effect. Consistent with prior work showing that disassembly of Golgi stacks accelerates protein trafficking (*26*), our findings therefore offer a mechanism for how histamine increases secretion of inflammatory factors.

Our study revealed that TG induces Golgi fragmentation through increasing cytosolic Ca^2+^ and GRASP55 phosphorylation. A similar case has been described previously in Alzheimer’s disease, where cytosolic calcium increase by Aβ treatment triggers activation of a cytosolic protease, calpain, which cleaves p35 to generate p25 and activate Cdk5, a cytoplasmic kinase that is highly expressed in neurons (*55*). Subsequently, activated Cdk5 phosphorylates GRASP65 and perhaps other Golgi structural proteins, leading to Golgi fragmentation (*13*). Although PKC and GRASP55 were not the focus in this study, expression of a phosphorylation deficient mutant of GRASP55 significantly reduced Golgi fragmentation as well as Aβ production. Taken together, our studies indicate that the Golgi is sensitive to cellular stimuli and stresses as in disease conditions, and responds to signaling cues to adjust its structure and function through increasing cytosolic Ca^2+^ and GRASP55 modification. Future studies defining the detailed mechanisms may help understand disease pathologies with Golgi and trafficking defects.

## MATERIALS AND METHODS

### Reagents, Plasmids and siRNA

All reagents used were from Sigma-Aldrich (St. Louis, MO), Roche (Basel, Switzerland) or Calbiochem (EMD Millipore, Burlington, MA), unless otherwise stated. The Annexin V apoptosis detection kit was from BioVision Inc. (San Francisco, CA). PKCα-GFP and GFP-PKCβII plasmids were provided by Dr. Yusuf Hannun (Stony Brook Cancer Center). GFP-PKCβI, GFP-PKCδ, and GFP-PKCζ plasmids were provided by Dr. Hesham El-Shewy (Medical University of South Carolina). The CAMKIIβ plasmid was provided by Dr. Mohammed Akaaboune (University of Michigan). The Str-li_VSVGwt-SBP-EGFP plasmid was provided by Dr. Franck Pere (Institut Curie). PKCγ-GFP cDNA construct was purchased from Addgene (Cambridge, MA). The HeLa cell line ManII-GFP was made in house and expresses α-mannosidase II covalently linked to green fluorescent protein.

Control siRNA (Silencer Select Negative Control #1 siRNA) was purchased from Applied Biosystems (ThermoFisher). PKC-specific custom siRNA targeting to endogenous human PKCα (5’-CAGAAGAACTGTATGCAAT-3’) was purchased from Ambien (ThermoFisher). To perform knockdowns, 200 nM of each oligo was used to transfect cells for 48 hours.

### Cell Culture and Drug Treatments

For all experiments, mycoplasma-free HeLa cells were obtained from ATCC (Manassas, VA) and passaged ≤20 times prior to use in experiments. HeLa cells were cultured in Dulbecco’s modified Eagle’s medium (DMEM; ThermoFisher, Waltham, MA) supplemented with 10% fetal bovine serum (FBS; Gemini Bio-Products, Sacramento, CA) and 100 units/ml penicillin-streptomycin at 37°C with 5% CO_2_. Cells were grown on glass coverslips according to standard tissue culture methods. Coverslips were pre-coated with poly-lysine (Gibco) to aid in cell attachment. For EGTA treatment, an excess of EGTA (8.96 mM) was combined with growth medium and applied to cells 1 h prior to TG treatment, as previously described (*56, 57*). All drugs, except cAMP-dependent protein kinase inhibitor (PKI) that is a peptide and dissolved in water, were made in DMSO, aliquoted, and stored at −20°C. Stocks were diluted into working solutions of DMEM at the time of the experiment as described in the text or figure legend. Upon the addition of the drug, cells were incubated at 5% CO_2_ and 37°C for the indicated times. Cells were washed 3 times with ice cold phosphate buffered saline (PBS) and collected with a cell scraper.

### Immunofluorescence Microscopy

For fluorescence microscopy, cells were rinsed 3 times in ice cold PBS, fixed with 4% (w/v) paraformaldehyde, quenched with 50 mM NH_4_Cl, permeabilized in 0.2% v/v Triton X-100 in PBS, and blocked for 1 h with PBS with 1% w/v bovine serum albumin (BSA) Fraction V (ThermoFisher, Waltham, MA) (*58*). Cells were incubated with a primary antibody diluted in 1% BSA in PBS (PBSB) at room temperature for 1.5 h, washed with PBS, and incubated with an FITC- or TRITC-labeled secondary antibody (1:200 dilution) in PBSB for 45 min at room temperature. Cells were washed 3 times with PBS and stained with 1:5,000 Hoechst dye for 5 min, mounted on glass slides with Moviol, and images were captured with a Zeiss (Oberkochen, Germany) Observer fluorescent microscope with a 63x oil objective lens with a numerical aperture of 1.4 and an Axiocam Mrm camera. TIF files were exported with AxioVision software (Zeiss).

For super-resolution microscopy, Alexa Fluor 647, and Alexa Fluor 488-labeled secondary antibodies (ThermoFisher) were used. After washing, coverslips were mounted to slides using ProLong Diamond antifade super resolution imaging mountant (ThermoFisher). Super-resolution images were imaged using Leica (Wetzlar, Germany) TCS SP8 STED super resolution microscope. Images were quantified using the NIH ImageJ software and assembled into figures with Photoshop Elements (Adobe, San Jose, CA). To clearly show the Golgi structure, brightness or contrast was adjusted linearly across all samples within each experiment.

To quantify Golgi fragmentation, cells were evaluated by eye under a microscope according to predefined fragmentation criteria, at least 300 cells were counted in each reaction. In experiments where transfected proteins were employed, only transfected cells were counted, and 100 cells were counted per replicate. Hoechst was used to identify individual, mitotic and overlapping cells. Briefly, cells having ≥3 disconnected Golgi items were considered fragmented, and cells with <3 were considered intact. In experiments where an inhibitor screen was performed, an unbiased image thresholding method was used to extract fragmentation data from ≥40 cells per replicate.

### Electron Microscopy (EM)

For EM, cells were fixed in pre-warmed serum-free DMEM, 20 mM HEPES, pH 7.4, 2% glutaraldehyde at room temperature for 30 min or 4°C overnight as previously described (*59*). Cells were washed 2 times with 0.1 M Sodium cacodylate (Electron Microscopy Sciences, Hatfield, PA), and post-fixed on ice in 1% v/v reduced Osmium tetroxide, 0.1 M Sodium cacodylate (w/v) and 1.5% cyanoferrate (w/v) in water. Cells were rinsed 3 times with 50 mM maleate buffer, pH 5.2, 3 times with water, scraped, and pelleted in microcentrifuge tubes for embedding. The EMBED 812 (EMD) protocol was used to embed cells and resin blocks were sectioned to 60 nm with a diamond knife and mounted on Formvar-coated copper grids. Samples were double contrasted with 2% uranyl acetate then with lead citrate and rinsed with copious amounts of water. Grids were imaged using a Philips (Amsterdam, Netherlands) transmission electron microscope. Golgi images were captured at 11,000x magnification. Golgi stacks were identified using morphological criteria and quantified using standard stereological techniques. A Golgi cisterna was identified as a perinuclear membrane within a Golgi stack ≥4 times longer than its width. Stack length was measured for the longest cisterna within a Golgi stack using the ruler tool in Photoshop Elements 13. For the number of cisternae per stack, the number of cisternae was counted. For the number of vesicles per stack, round objects no greater than 80 µm in diameter within 0.5 µm of a Golgi stack were counted. At least 20 cells were quantified in each experiment, and the EM results represent two independent experiments.

### Protein Biochemistry and Antibodies

For immunoblotting, cells from a 10 cm dish were pelleted and lysed with 30 µl lysis buffer (20 nM Tris-HCl, 150 mM NaCl, 1% Triton X-100 (Bio-Rad, Hercules, CA), 20 mM beta-glycerol phosphate and protease inhibitors cocktail, NaF, NaVan, pH 8.0). Samples were made with 6X SDS PAGE buffer with fresh DTT, denatured at 95°C for 4 min and then run on PAGE gels. Protein was transferred to nitrocellulose membranes using semi-dry transfer at constant 16 V. Membranes were blocked for 10 min with 3% milk in 0.2% Tween-20 in phosphate buffered saline (PBST) and immunoblotted. The following antibodies were used: monoclonal antibodies against β-actin and GFP (Sigma-Aldrich), Gos28 and GM130 (BD Biosciences, Franklin Lanes, NJ), PKCα (Santa Cruz Biotechnology, Dallas, TX), and α-tubulin (Developmental Studies Hybridoma Bank, University of Iowa); polyclonal antibodies against CHOP, p-eIF2α, eIF2α and p115 (Cell Signaling, Danvers, MA), GRASP55 and GRASP65 (Proteintech), Bip (Santa Cruz), GM130 (“N73” from J. Seemann), and TGN46 (Bio-Rad). 1:200 goat anti-mouse, 1:200 goat anti-sheep (for TGN46) and 1:5,000 goat anti-rabbit HRP-conjugated secondary antibodies were used. Western blots were captured with Enhanced Chemiluminescence (ECL) dye reagent (ThermoFisher), in a FluorChem M chemi-luminescent imager (ProteinSimple, San Jose, CA).

### VSV-G trafficking using RUSH system

VSV-G trafficking was performed as previously described (*41*). Briefly, HeLa cells were transfected with the Str-li_VSVG wt-SBP-EGFP plasmid (*40*) and cultured at 37°C for 16 h. Cells were then incubated with 250 nM TG or 10 µM monensin in fresh medium for 0.5 h at 37°C before 40 µM D-biotin (VWR Life Science, Radnor, PA) was added. Cells were then lysed at indicated time points (chase), treated with or without EndoH, and analyzed by Western blotting for VSV-G-GFP using a GFP antibody. The percentage of EndoH resistant VSV-G was quantified using the ImageJ software.

### Molecular Cloning

Constructs for GRASP55 truncation mutants, aa1-212, aa1-250, aa1-300, aa1-400, aa1-430 were constructed in pEGFP-N1 vector using BamHI and HindIII sites (*47*). All cDNAs generated in this study were confirmed by DNA sequencing.

### Quantitation and Statistics

All data represent the mean ± SEM (standard error of the mean) of at least three independent experiments unless noted. A statistical analysis was conducted with two-tailed Student’s t-test in the Excel program (Microsoft, Redmond, WA). Differences in means were considered statistically significant at p < 0.05. Significance levels are: *, p<0.05; **, p<0.01; ***, p<0.001. Figures were assembled with Photoshop (Adobe, San Jose, CA).

## AUTHOR CONTRIBUTIONS

Conception and design: S. Ramnarayanan, S. Ireland, Y. Wang

Development of methodology: S. Ireland, S. Ramnarayanan, M. Fu, X. Zhang, D. Emebo, Y. Wang

Acquisition of data: S. Ireland, S. Ramnarayanan, M. Fu, X. Zhang, D. Emebo

Analysis and interpretation of data: S. Ireland, S. Ramnarayanan, M. Fu, Y. Wang

Writing, review, and/or revision of the manuscript: S. Ireland, Y. Wang

Administrative, technical, or material support: Y. Wang, S. Ireland

Study supervision: Y. Wang

## AKNOWLEDGMENTS

We thank Drs. Yusuf Hannun, Hesham El-Shewy, Isabel Martinez Peña, Mohammed Akaaboune and Franck Perez for reagents; Drs. Peter Blumberg, John Kuwada, Edward Stuenkel, Haoxing Xu for technical assistance; Gregg Sobocinski for microscope and equipment expertise; Dana Holcomb for microscopy help. This work was supported by the National Institutes of Health (Grants GM112786, GM105920, and GM130331), MCubed and the Fastforward Protein Folding Disease Initiative of the University of Michigan to Y. Wang. S. Ireland is a graduate student who has been partially supported by the Mary Sue and Kenneth Coleman Endowed Fellowship Fund from the University of Michigan. Many thanks to past and current members of the Wang lab, including Erpan Ahat, Michael Bekier, Shijiao Huang, Courtney Killeen, Haoran Huang, and Ron Benyair.

## ABBREVIATIONS

B/AM: BAPTA-AM
CAMKII: Ca^2+^/calmodulin-dependent protein kinase II
DTT: dithiothreitol
EM: electron microscopy
ER: endoplasmic reticulum
Endo H: Endoglycosidase H
GFP: green fluorescent protein
GORASP1: Golgi reassembly stacking protein 1
GORASP2: Golgi reassembly stacking protein 2
GRASP55: Golgi reassembly stacking protein of 55 kDa
GRASP65: Golgi reassembly stacking protein of 65 kDa
PBS: phosphate-buffered saline
PEI: polyethylenimine
PFA: paraformaldehyde
SPR domain: serine/proline-rich domain
TGN: *trans*-Golgi network
PKA: Protein Kinase A
PKC: Protein Kinase C
SNARE: SNAP receptor protein
PMA: phorbol 12-myristate 13-acetate
TG: thapsigargin
TGN: *trans*-Golgi network
Tu: tunicamycin
VSV-G: vesicular stomatitis virus glycoprotein

## Supplementary Figures

**Figure S1.**
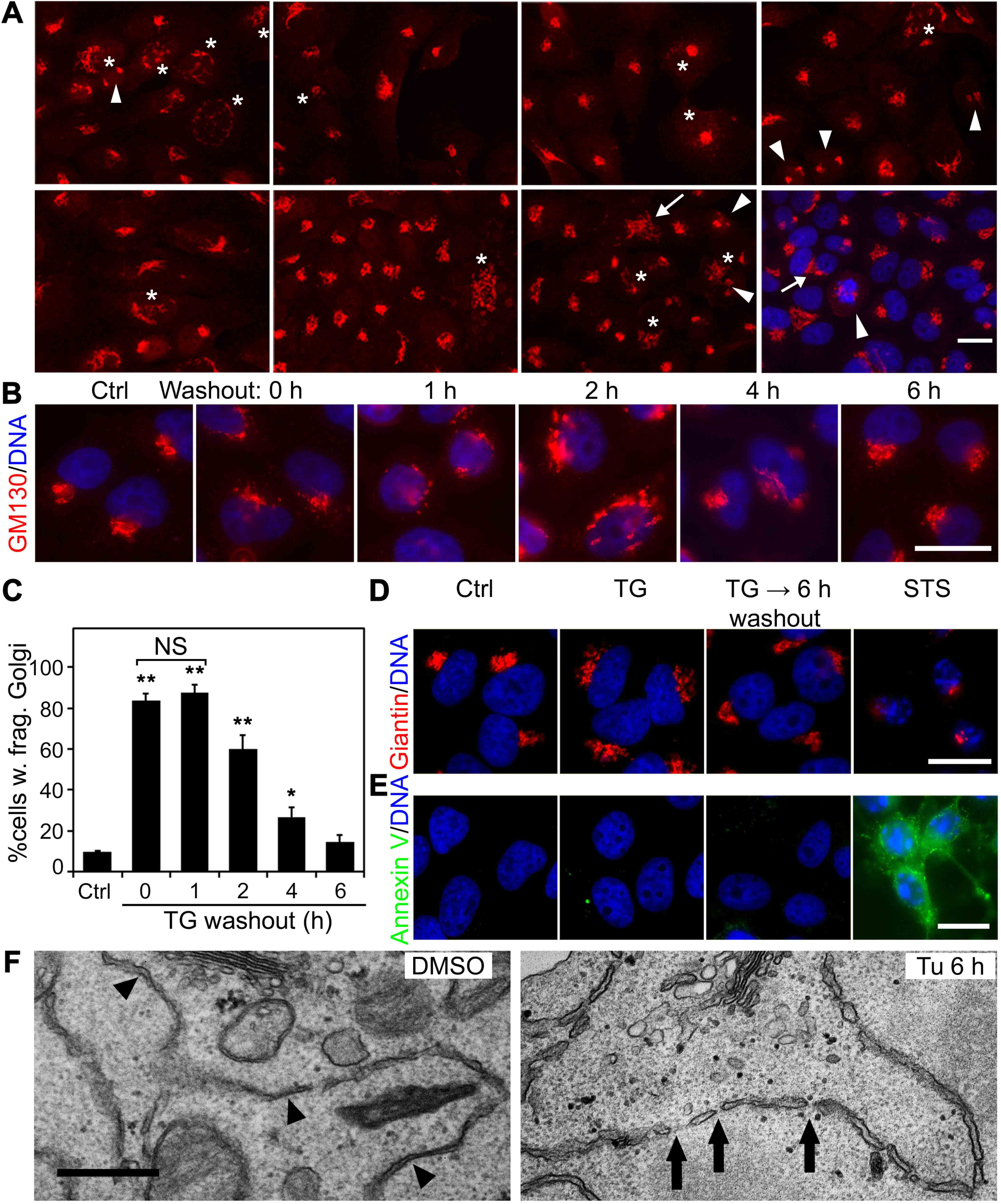
TG-induced Golgi fragmentation is reversible. **(A)** A gallery of cells with intact or fragmented Golgi. Random images of HeLa cells treated with or without TG were labeled with a GM130 antibody. Asterisks (*) indicate fragmented Golgi, arrowheads (►) are mitotic Golgi, arrows (**→)** are overlapping Golgi from two or more distinct cells. The last frame includes the DNA channel in blue to show a mitotic cell. The following criteria are used to define whether a Golgi is intact or fragmented: 1) If the Golgi exists as a single piece of connected membrane, it is intact. 2) If a Golgi exhibits several items that are connected by visible membrane bridges, even though these bridges might be faint, the Golgi is considered intact. 3) If a Golgi exhibits ≥ 3 disconnected pieces (no visible bridges connecting them), then the Golgi is fragmented. 4) Mitotic cells, defined by the DNA pattern, and overlapping cells in which the Golgi pattern is difficult to define, are not counted. **(B)** TG-induced Golgi fragmentation is reversible. Cells were treated with either DMSO or 100 nM TG for 1 h. After washing out TG, TG-treated cells were incubated in fresh growth medium for the indicated times and stained for GM130 and DNA. **(C)** Quantitation of (B) for cells with fragmented Golgi. For statistical analyses, treated cells were compared to the DMSO control (Ctrl). *, p ≤ 0.05; **, p ≤ 0.01. **(D)** Acute TG treatment does not cause apoptosis. Cells were treated with either DMSO or 100 nM TG for 20 min without or with 6 h washout, or with 2 µM staurosporine (STS) for 4 h, and stained for GM130 and DNA. **(E)** Cells in (D) were surface stained with Annexin V-EGFP. Scale bars in all fluorescent images, 20 µm. **(F)** Representative EM images of ER cisternae in cells treated with DMSO (Ctrl) or tunicamycin (Tu, 5 µg/ml) for 6 h. ER cisternae in Ctrl cells, indicated by arrowheads (►), appear to have a narrow, intact structure; where in Tu treated cells, the ER cisternae appear to be swollen and fragmented, as indicated by arrows (**→**). Scale bar, 0.5 µm.

**Figure S2.**
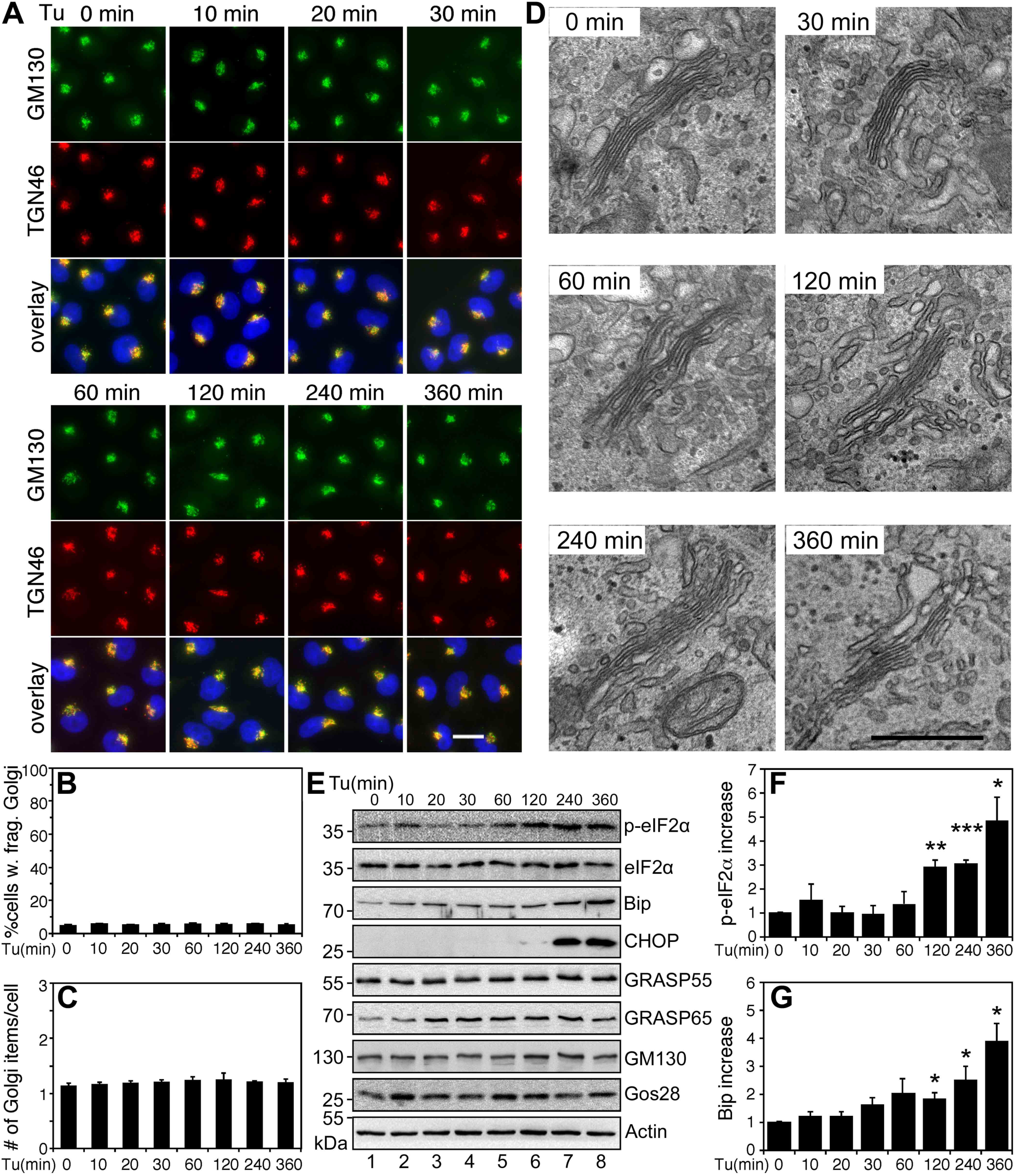
Tu treatment induces UPR but not Golgi fragmentation. **(A)** Tu treatment has no effect on the Golgi morphology. HeLa cells were treated with 5 µg/ml Tu for indicated times and stained for GM130 and TGN46. Scale bar, 20 µm. **(B-C)** Quantitation of Golgi fragmentation in Tu-treated cells in (A). **(D)** EM micrographs of the Golgi region in Tu-treated cells. Scale bar, 0.5 µm. **(E)** Tu-treated cells as in (A) were analyzed by Western blots. Note the increased levels of p-eIF2*α*, Bip and CHOP over time. **(F-G)** Quantitation of p-eIF2α /eIF2α and Bip in (E). Results are shown as Mean ± SEM from at least 3 independent experiments; statistical analyses were performed using two-tailed Student’s *t*-tests (*, p ≤ 0.05; **, p ≤ 0.01; ***, p ≤ 0.001).

**Figure S3.**
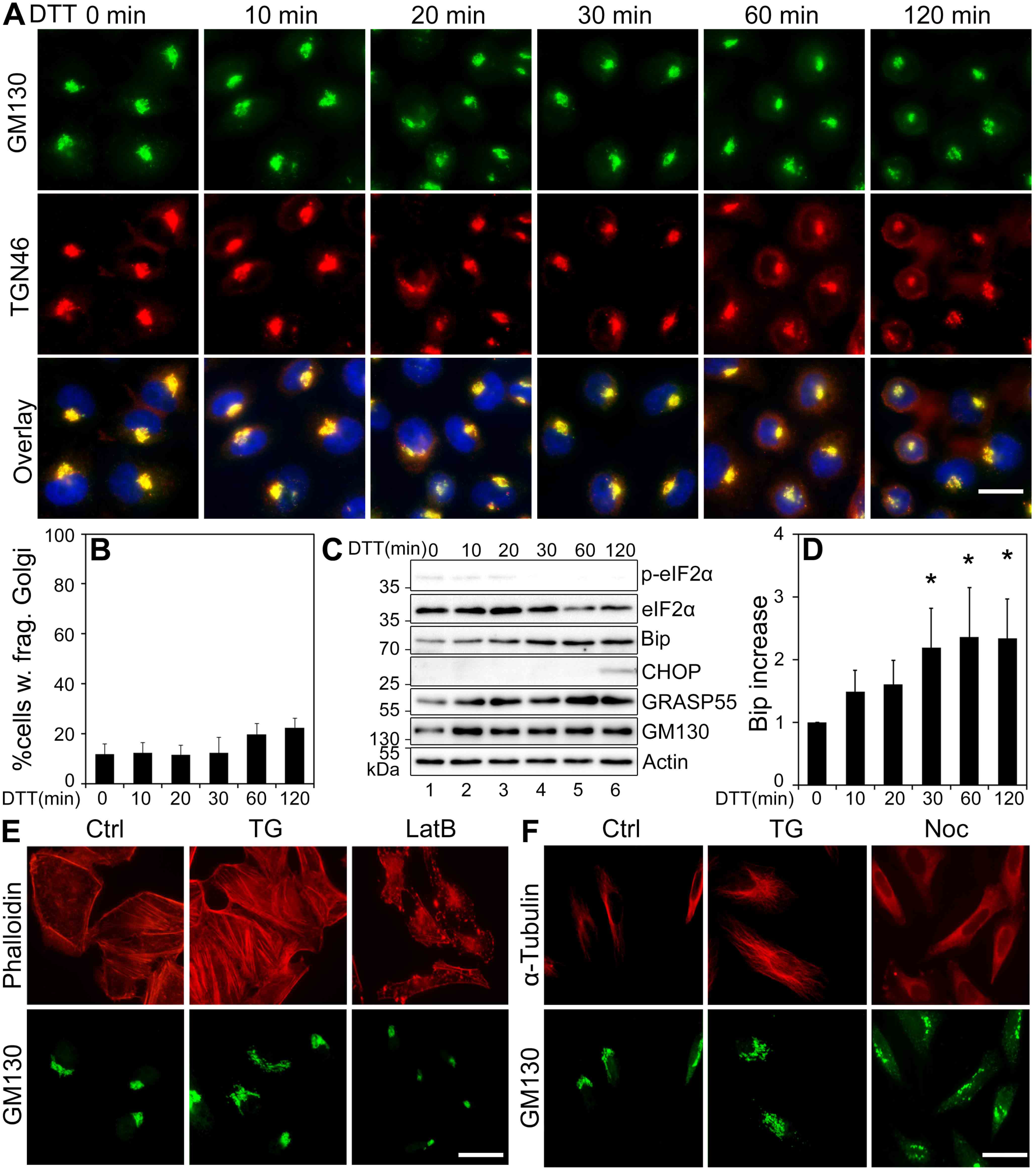
DTT-treatment induces UPR but not Golgi fragmentation. **(A)** HeLa cells were treated with DMSO or 10 mM DTT for indicated times and stained for GM130 (green), and TGN46 (red). Scale bar, 20 µm. **(B)** Quantitation of (A) for cells with fragmented Golgi. **(C)** Western blots of ER stress and Golgi proteins showing UPR induction upon DTT treatment. **(D)** Quantitation of Bip levels from four independent experiments. Two-tailed Student’s *t*-tests were used to calculate statistical significance (*, p ≤ 0.05). **(E)** Cells were treated with 0.5 µM latrunculin B for 2 h and 250 nM TG was added for the last 20 min. Cells were stained for F-actin with phalloidin (red) and GM130 (green). Scale bar, 20 µm. **(F)** Cells were treated with 1 µM Noc for 2 h and 250 nM TG was added for the last 20 min. Cells were stained for α-tubulin (red) and GM130 (green). Scale bar, 20 µm.

**Figure S4.**
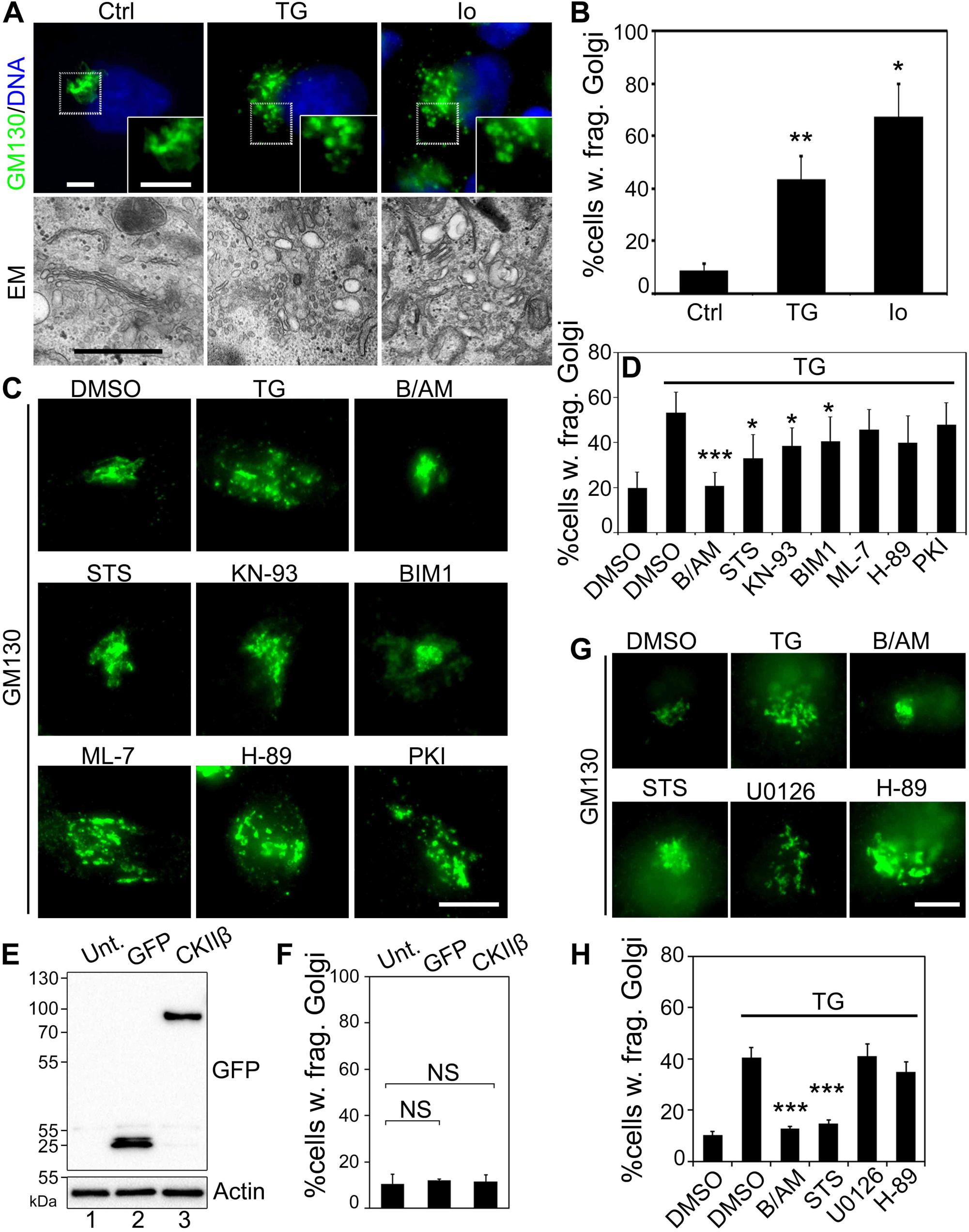
Identification of PKC as the kinase for TG-induced Golgi fragmentation. **(A)** HeLa cells were treated with DMSO (Ctrl), 100 nM TG, or 1 µM ionomycin (Io) for 1 h, and analyzed by fluorescence microscopy (top panels) and EM (lower panels). Scale bars on the fluorescence images, 5 µm; scale bar on EM micrographs, 0.5 µm. **(B)** Quantitation of cells with fragmented Golgi in (A) based on the GM130 pattern. **(C)** PKC inhibition reduces TG-induced Golgi fragmentation. Cells were pre-treated with 60 µM BAPTA-AM (B/AM), 2 µM staurosporine (STS), 5 µM KN-93, 2 µM BIM1, 5 µM ML-7, 30 µM H-89, or 10 µM PKI for 10 min, followed by the treatment with either DMSO or 250 nM TG for 20 min, and stained for GM130. Scale bar, 10 µm. **(D)** Quantitation of (C) for Golgi fragmentation. Experiments were quantified in a double-blinded fashion and results from three independent experiments were used to calculate means ± SEM. **(E)** Western blot of cells transfected with GFP or GFP-CAMKIIβ using a GFP antibody. **(F)** Quantitation of cells with fragmented Golgi in cells transfected with GFP or GFP-CAMKIIβ as in (E). For statistics, transfected cells were compared to control untransfected (Unt.) cells. **(G)** HeLa cells were treated with MAPK/ERK or PKA/PKD inhibitors, U0126 and H-89, respectively, followed by 100 nM TG treatment for 20 min. **(H)** Quantitation of Golgi fragmentation in (G). Two-tailed Student’s *t*-tests were used to calculate statistical significance (*, p ≤ 0.05; **, p ≤ 0.01; ***, p ≤ 0.001).

**Figure S5.**
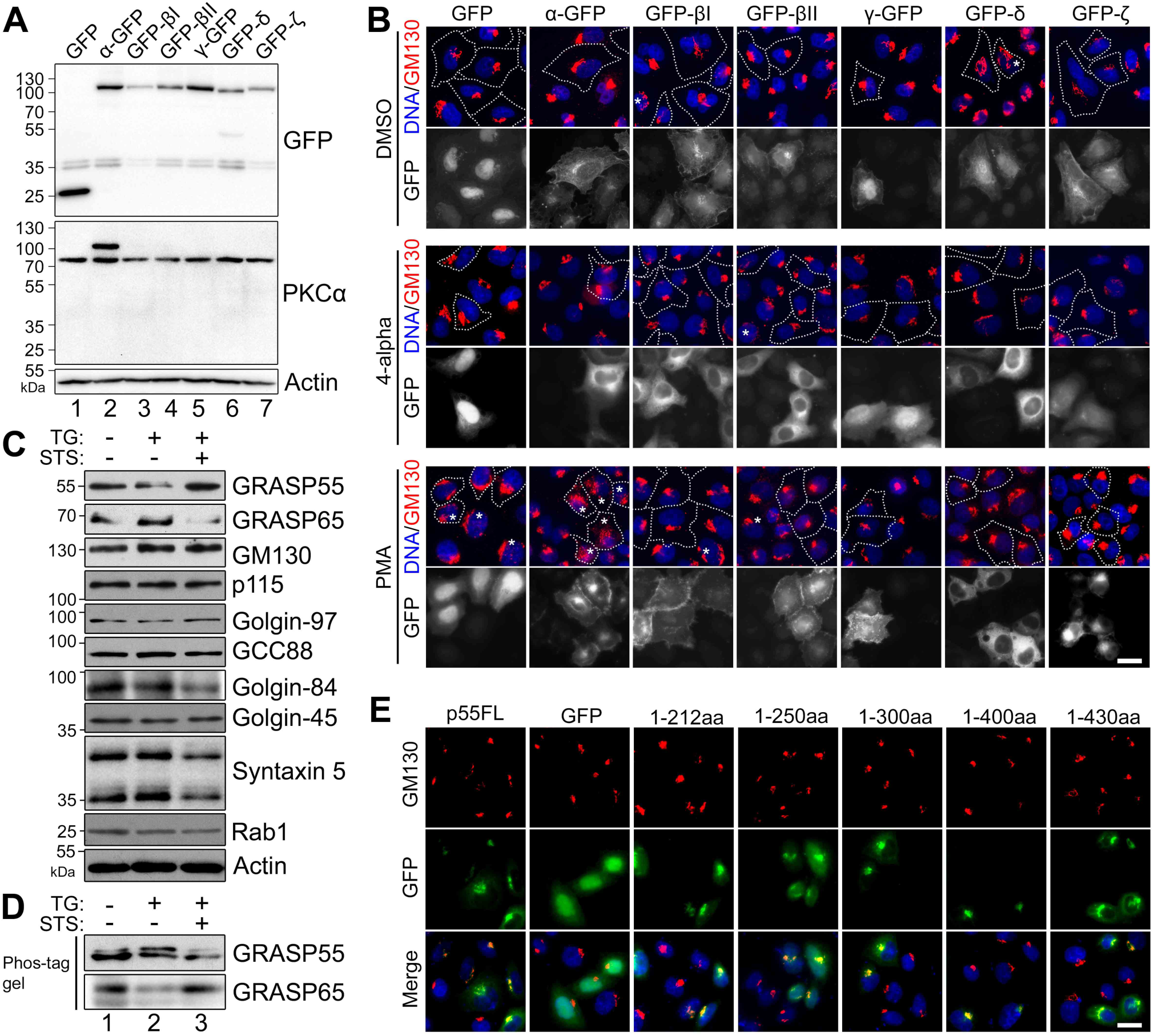
PKCα localizes to the Golgi upon activation with PMA. **(A)** Expression of PKC isoforms. HeLa cells were transfected with indicated PKC isoforms and analyzed by Western blot for GFP or PKCα. **(B)** Fluorescent images showing the localization of expressed PKC isoforms after treatment with DMSO, 4-alpha, or PMA. PMA induces PKCα (α-GFP) localization to the Golgi area, a similar but less dramatic effect was observed for PKCβII (GFP-βII). Scale bar, 20 µm. **(C)** TG treatment results in the phosphorylation of GRASP55 but not other Golgi proteins. HeLa cells were treated with 250 nM TG for 1 h, with or without 2 µM staurosporine (STS) pre-treatment for 10 min, and analyzed by Western blot. **(D)** Cell lysates in (C) were analyzed by Phos-tag gels and Western blot. **(E)** Immunofluorescence images of cells expressing indicated GRASP55 truncation mutants.

**Figure S6.**
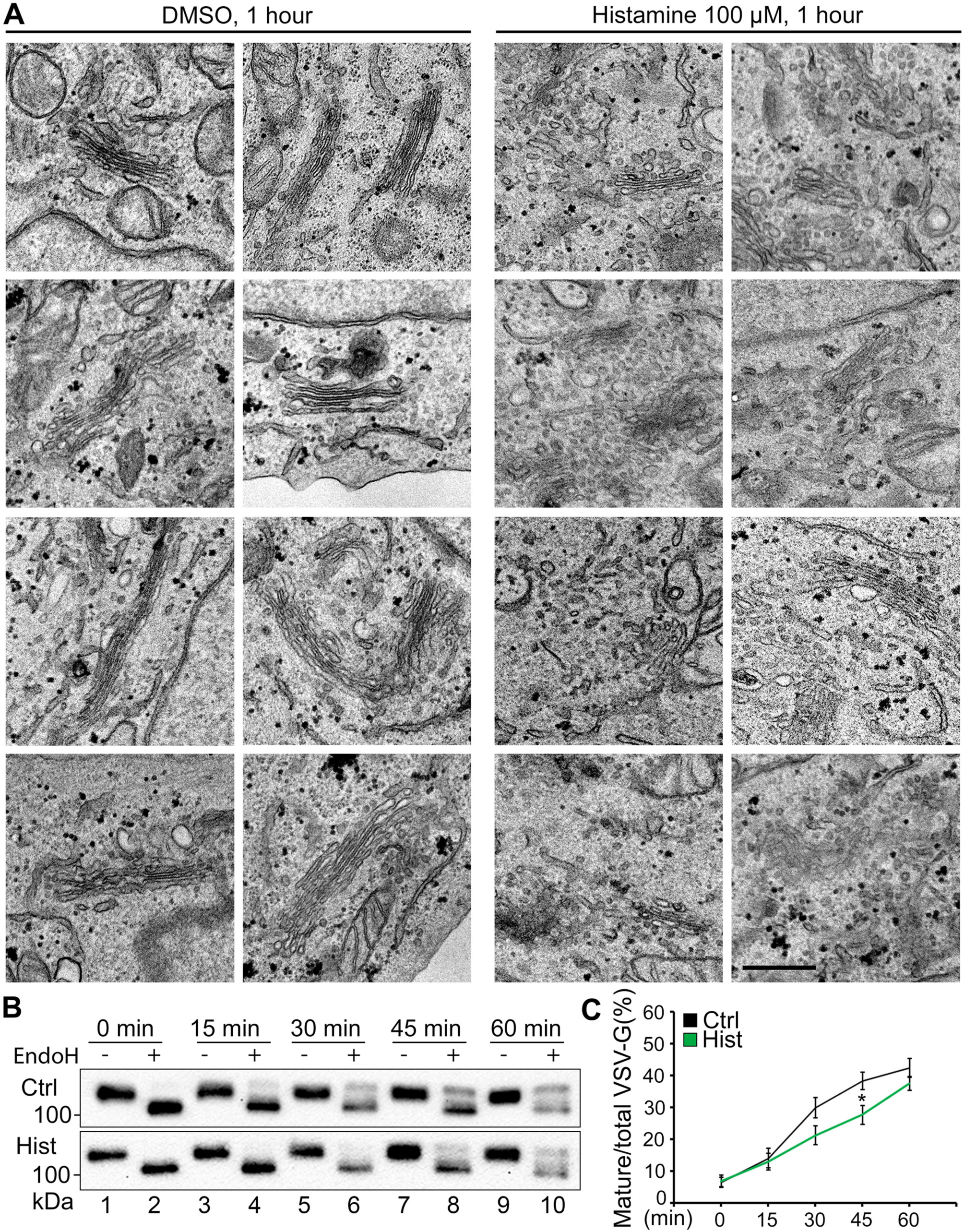
Histamine treatment alters the Golgi structure. **(A)** HeLa cells treated with either DMSO (control, 1 h) or histamine (100 µM, 1 h) were analyzed by EM. Shown are collections of electron micrographs representing the two treatments. Consistent aberrations in Golgi shape were frequently seen in histamine-treated cells, including reduced cisternae number and stack length, and increased number of vesicles. Scale bar, 0.5 µm. **(B)** Cells were transfected with the Str-li_VSVG wt-SBP-EGFP plasmid for 16 h and treated without (Ctrl) or with 100 µM histamine (Hist) at 37°C for 1 h. Cells were then incubated with complete medium containing 40 µM biotin and 100 µM of Hist, lysed at the indicated time points, and treated without (-) or with (+) EndoH, and analyzed by Western blot. **(C)** Quantification of (B) for the percentage of EndoH resistant VSV-G from 3 independent experiments. All quantitation results are shown as Mean ± SEM. Statistical analyses were performed using two-tailed Student’s *t*-tests (*, p ≤ 0.05).

## REFERENCES

1. M. S. Ladinsky, D. N. Mastronarde, J. R. McIntosh, K. E. Howell, L. A. Staehelin, Golgi structure in three dimensions: functional insights from the normal rat kidney cell. J Cell Biol 144, 1135–1149 (1999).

2. Y. Wang, J. Seemann, M. Pypaert, J. Shorter, G. Warren, A direct role for GRASP65 as a mitotically regulated Golgi stacking factor. The EMBO journal 22, 3279–3290 (2003).

3. X. Zhang, Y. Wang, GRASPs in Golgi Structure and Function. Frontiers in Cell and Developmental Biology 3, 84 (2015).

4. Y. Wang, J. Seemann, Golgi biogenesis. Cold Spring Harb Perspect Biol 3, a005330 (2011).

5. Y. Xiang, Y. Wang, GRASP55 and GRASP65 play complementary and essential roles in Golgi cisternal stacking. J Cell Biol 188, 237–251 (2010).

6. M. A. Puthenveedu, C. Bachert, S. Puri, F. Lanni, A. D. Linstedt, GM130 and GRASP65-dependent lateral cisternal fusion allows uniform Golgi-enzyme distribution. Nature cell biology 8, 238–248 (2006).

7. T. N. Feinstein, A. D. Linstedt, GRASP55 Regulates Golgi Ribbon Formation. Molecular biology of the cell 19, 2696–2707 (2008).

8. T. Jarvela, A. D. Linstedt, Isoform-specific tethering links the Golgi ribbon to maintain compartmentalization. Molecular biology of the cell, (2013).

9. Y. Wang, A. Satoh, G. Warren, Mapping the functional domains of the Golgi stacking factor GRASP65. J Biol Chem 280, 4921–4928 (2005).

10. W. J. Krause, Brunner’s Glands: A Structural,Histochemical and Pathological Profile. Progress in Histochemistry and Cytochemistry 35, 255–367 (2000).

11. D. A. Thayer, Y. N. Jan, L. Y. Jan, Increased neuronal activity fragments the Golgi complex. Proc Natl Acad Sci U S A 110, 1482–1487 (2013).

12. A. Rambourg, Y. Clermont, M. Chretien, L. Olivier, Modulation of the Golgi apparatus in stimulated and nonstimulated prolactin cells of female rats. Anat Rec 235, 353–362 (1993).

13. G. Joshi, M. E. Bekier, 2nd, Y. Wang, Golgi fragmentation in Alzheimer’s disease. Front Neurosci 9, 340 (2015).

14. Y. Mizuno et al., Familial Parkinson’s disease. Alpha-synuclein and parkin. Adv Neurol 86, 13–21 (2001).

15. P. Hilditch-Maguire et al., Huntingtin: an iron-regulated protein essential for normal nuclear and perinuclear organelles. Hum Mol Genet 9, 2789–2797 (2000).

16. Y. Fujita, K. Okamoto, Golgi apparatus of the motor neurons in patients with amyotrophic lateral sclerosis and in mice models of amyotrophic lateral sclerosis. Neuropathology 25, 388–394 (2005).

17. N. K. Gonatas, J. O. Gonatas, A. Stieber, The involvement of the Golgi apparatus in the pathogenesis of amyotrophic lateral sclerosis, Alzheimer’s disease, and ricin intoxication. Histochemistry and cell biology 109, 591–600 (1998).

18. Z. Mourelatos, N. K. Gonatas, A. Stieber, M. E. Gurney, M. C. Dal Canto, The Golgi apparatus of spinal cord motor neurons in transgenic mice expressing mutant Cu,Zn superoxide dismutase becomes fragmented in early, preclinical stages of the disease. Proc Natl Acad Sci U S A 93, 5472–5477 (1996).

19. X. Tan et al., Epithelial-to-mesenchymal transition drives a pro-metastatic Golgi compaction process through scaffolding protein PAQR11. J Clin Invest, (2016).

20. A. Petrosyan, M. S. Holzapfel, D. E. Muirhead, P. W. Cheng, Restoration of compact Golgi morphology in advanced prostate cancer enhances susceptibility to galectin-1-induced apoptosis by modifying mucin O-glycan synthesis. Mol Cancer Res 12, 1704–1716 (2014).

21. R. Sewell et al., The ST6GalNAc-I sialyltransferase localizes throughout the Golgi and Is responsible for the synthesis of the tumor-associated sialyl-Tn O-glycan in human breast cancer. J Biol Chem 281, 3586–3594 (2006).

22. C. Sutterlin, R. Polishchuk, M. Pecot, V. Malhotra, The Golgi-associated protein GRASP65 regulates spindle dynamics and is essential for cell division. Molecular biology of the cell 16, 3211–3222 (2005).

23. D. Tang, H. Yuan, Y. Wang, The Role of GRASP65 in Golgi Cisternal Stacking and Cell Cycle Progression. Traffic 11, 827–842 (2010).

24. M. E. Bekier, 2nd et al., Knockout of the Golgi stacking proteins GRASP55 and GRASP65 impairs Golgi structure and function. Molecular biology of the cell 28, 2833–2842 (2017).

25. E. Ahat, J. Li, Y. Wang, New Insights Into the Golgi Stacking Proteins. Frontiers in Cell and Developmental Biology In press, (2019).

26. Y. Xiang et al., Regulation of protein glycosylation and sorting by the Golgi matrix proteins GRASP55/65. Nat Commun 4, 1659 (2013).

27. E. Ahat, Y. Xiang, X. Zhang, M. E. Bekier, Y. Wang, GRASP depletion-mediated Golgi destruction decreases cell adhesion and migration via the reduction of alpha5beta1 integrin. Molecular biology of the cell 30, 766–777 (2019).

28. X. Zhang et al., GORASP2/GRASP55 collaborates with the PtdIns3K UVRAG complex to facilitate autophagosome-lysosome fusion. Autophagy, 1–14 (2019).

29. B. S. E. D. El Homasany et al., The Scaffolding Protein CG-NAP/AKAP450 Is a Critical Integrating Component of the LFA-1-Induced Signaling Complex in Migratory T Cells. The Journal of Immunology 175, 7811–7818 (2005).

30. D. Chen, A. Purohit, E. Halilovic, S. J. Doxsey, A. C. Newton, Centrosomal anchoring of protein kinase C betaII by pericentrin controls microtubule organization, spindle function, and cytokinesis. J Biol Chem 279, 4829–4839 (2004).

31. Y. Nishizuka, Protein kinase C and lipid signaling for sustained cellular responses. FASEB J 9, 484–496 (1995).

32. H. Farhan et al., MAPK signaling to the early secretory pathway revealed by kinase/phosphatase functional screening. J Cell Biol 189, 997–1011 (2010).

33. M. Cooke, A. Magimaidas, V. Casado-Medrano, M. G. Kazanietz, Protein kinase C in cancer: The top five unanswered questions. Mol Carcinog 56, 1531–1542 (2017).

34. H. Kim, R. Zamel, X. H. Bai, M. Liu, PKC activation induces inflammatory response and cell death in human bronchial epithelial cells. PLoS One 8, e64182 (2013).

35. E. M. Griner, M. G. Kazanietz, Protein kinase C and other diacylglycerol effectors in cancer. Nat Rev Cancer 7, 281–294 (2007).

36. M. Oku et al., Novel cis-acting element GASE regulates transcriptional induction by the Golgi stress response. Cell Struct Funct 36, 1–12 (2011).

37. C. Xu, H. Ma, G. Inesi, M. K. Al-Shawi, C. Toyoshima, Specific structural requirements for the inhibitory effect of thapsigargin on the Ca2+ ATPase SERCA. J Biol Chem 279, 17973–17979 (2004).

38. Y. Ito, Y. Takeda, A. Seko, M. Izumi, Y. Kajihara, Functional analysis of endoplasmic reticulum glucosyltransferase (UGGT): Synthetic chemistry’s initiative in glycobiology. Semin Cell Dev Biol, (2014).

39. K. T. Jones, G. R. Sharpe, Thapsigargin raises intracellular free calcium levels in human keratinocytes and inhibits the coordinated expression of differentiation markers. Exp Cell Res 210, 71–76 (1994).

40. G. Boncompain et al., Synchronization of secretory protein traffic in populations of cells. Nat Methods 9, 493–498 (2012).

41. J. Li, D. Tang, S. C. Ireland, Y. Wang, DjA1 maintains Golgi integrity via interaction with GRASP65. Molecular biology of the cell 30, 478–490 (2019).

42. S. J. Fliesler, S. F. Basinger, Monensin stimulates glycerolipid incorporation into rod outer segment membranes. J Biol Chem 262, 17516–17523 (1987).

43. G. Joshi, Y. Chi, Z. Huang, Y. Wang, Abeta-induced Golgi fragmentation in Alzheimer’s disease enhances Abeta production. Proc Natl Acad Sci U S A 111, E1230–1239 (2014).

44. S. A. Jesch, T. S. Lewis, N. G. Ahn, A. D. Linstedt, Mitotic phosphorylation of Golgi reassembly stacking protein 55 by mitogen-activated protein kinase ERK2. Molecular biology of the cell 12, 1811–1817 (2001).

45. C. Jamora et al., Gbetagamma-mediated regulation of Golgi organization is through the direct activation of protein kinase D. Cell 98, 59–68 (1999).

46. T. Kajimoto, S. Ohmori, Y. Shirai, N. Sakai, N. Saito, Subtype-specific translocation of the delta subtype of protein kinase C and its activation by tyrosine phosphorylation induced by ceramide in HeLa cells. Molecular and cellular biology 21, 1769–1783 (2001).

47. X. Zhang et al., GRASP55 Senses Glucose Deprivation through O-GlcNAcylation to Promote Autophagosome-Lysosome Fusion. Developmental cell 45, 245–261 e246 (2018).

48. M. Matsubara, T. Tamura, K. Ohmori, K. Hasegawa, Histamine H1 receptor antagonist blocks histamine-induced proinflammatory cytokine production through inhibition of Ca2+-dependent protein kinase C, Raf/MEK/ERK and IKK/I kappa B/NF-kappa B signal cascades. Biochem Pharmacol 69, 433–449 (2005).

49. D. K. Saini et al., Regulation of Golgi structure and secretion by receptor-induced G protein betagamma complex translocation. Proc Natl Acad Sci U S A 107, 11417–11422 (2010).

50. N. Sahoo et al., Gastric Acid Secretion from Parietal Cells Is Mediated by a Ca(2+) Efflux Channel in the Tubulovesicle. Developmental cell 41, 262–273 e266 (2017).

51. G. Xie et al., Modulation of Mast Cell Toll-Like Receptor 3 Expression and Cytokines Release by Histamine. Cell Physiol Biochem 46, 2401–2411 (2018).

52. H. Y. Gee, S. H. Noh, B. L. Tang, K. H. Kim, M. G. Lee, Rescue of DeltaF508-CFTR Trafficking via a GRASP-Dependent Unconventional Secretion Pathway. Cell 146, 746–760 (2011).

53. A. Karpati et al., Histamine elicits glutamate release from cultured astrocytes. Journal of pharmacological sciences, (2018).

54. S. Sohen et al., Activation of histamine H1 receptor results in enhanced proteoglycan synthesis by human articular chondrocyte: involvement of protein kinase C and intracellular Ca(2+). Pathophysiology 8, 93–98 (2001).

55. J. Lew et al., A brain-specific activator of cyclin-dependent kinase 5. Nature 371, 423–426 (1994).

56. Q. Jiang et al., Increase of cytosolic calcium induced by trichosanthin suppresses cAMP/PKC levels through the inhibition of adenylyl cyclase activity in HeLa cells. Mol Biol Rep 38, 2863–2868 (2011).

57. M. Z. Cui, G. C. Parry, T. S. Edgington, N. Mackman, Regulation of tissue factor gene expression in epithelial cells. Induction by serum and phorbol 12-myristate 13-acetate. Arterioscler Thromb 14, 807–814 (1994).

58. D. Tang et al., Mena-GRASP65 interaction couples actin polymerization to Golgi ribbon linking. Molecular biology of the cell 27, 137–152 (2016).

59. D. Tang, Y. Xiang, Y. Wang, Reconstitution of the cell cycle-regulated Golgi disassembly and reassembly in a cell-free system. Nat Protoc 5, 758–772 (2010).

